# Distinct dopaminergic spike-timing-dependent plasticity rules are suited to different functional roles

**DOI:** 10.1101/2024.06.24.600372

**Authors:** Baram Sosis, Jonathan E. Rubin

**Affiliations:** Department of Mathematics, University of Pittsburgh, 301 Thackeray Hall, Pittsburgh, 15260, PA, USA; Berkeley, CA, USA; Center for the Neural Basis of Cognition, University of Pittsburgh, 4400 Fifth Ave, Pittsburgh, 15213, PA, USA

**Keywords:** Dopamine, Synaptic plasticity, STDP, Basal ganglia, Reward prediction, Value estimation, Action selection

## Abstract

Various mathematical models have been formulated to describe the changes in synaptic strengths resulting from spike-timing-dependent plasticity (STDP). A subset of these models include a third factor, dopamine, which interacts with spike timing to contribute to plasticity at specific synapses, notably those from cortex to striatum at the input layer of the basal ganglia. Theoretical work to analyze these plasticity models has largely focused on abstract issues, such as the conditions under which they may promote synchronization and the weight distributions induced by inputs with simple correlation structures, rather than on scenarios associated with specific tasks, and has generally not considered dopamine-dependent forms of STDP. In this paper we introduce forms of dopamine-modulated STDP adapted from previously proposed plasticity rules. We then analyze, mathematically and with simulations, their performance in two biologically relevant scenarios. We test the ability of each of the three models to complete simple value estimation and action selection tasks, studying the learned weight distributions and corresponding task performance in each setting. Interestingly, we find that each plasticity rule is well suited to a subset of the scenarios studied but falls short in others. Different tasks may therefore require different forms of synaptic plasticity, yielding the prediction that the precise form of the STDP mechanism present may vary across regions of the striatum, and other brain areas impacted by dopamine, that are involved in distinct computational functions.

## 1 Introduction

Learning and memory are critical features of cognition and a number of neural learning mechanisms have been described. One important mechanism is spike-timing-dependent plasticity (STDP) [1, 2], a class of Hebbian plasticity rules in which the relative timing of pre- and postsynaptic spikes determines the changes in synaptic connection strength. Typically, a presynaptic spike before a postsynaptic spike – that is, a causal ordering of the spikes – leads to synaptic potentiation, whereas the reverse order leads to depression. In some cases, however, synaptic plasticity depends not just on the timing of pre- and postsynaptic spikes but also on some third factor, such as a neuromodulatory signal or other input [3, 4]. These additional factors may act as gating signals, and their strength and timing may impact both the magnitude and the direction of synaptic changes.

A prominent example of neuromodulatory impact on synaptic plasticity occurs at the cortical inputs to the basal ganglia. The neuromodulator dopamine is released by midbrain dopamine neurons when unexpected reward is received [5, 6] and plays a crucial role in modulating plasticity of corticostriatal synapses [7]. Experimental evidence and theoretical modeling suggest that dopamine serves as a reward prediction error signal [8, 9], enabling the brain to learn to favor behaviors that lead to reward and disfavoring behaviors that do not; such findings have been recently reviewed [10]. Theoretical analysis of action selection and modulation by cortico-basal ganglia-thalamic (CBGT) circuits posits a role for these dopaminergic reward prediction error signals both in updating value estimates associated with available choices and, through their impact on corticostriatal synaptic strengths, in altering the likelihood that a particular action will be selected in the future [11–14]. These distinct functions are likely performed by different neurons in different regions of the basal ganglia, however, which raises the possibility that distinct plasticity rules are involved. Unfortunately, despite some exciting experimental investigations of long-term plasticity properties in specific striatal regions and task settings [15–17], relatively little is known about the details of these plasticity mechanisms, especially in ventral striatal regions thought to encode value, which project to dopaminergic areas in the substantia nigra and hence are well positioned to contribute to calculation of reward prediction error [18, 19].

Previous studies [14, 20, 21] have introduced computational models of reinforcement learning and action selection in the basal ganglia in which the strengths of the corticostriatal synapses determine the likelihood that each action will be selected. While these simulations employ biologically plausible neural models and synaptic plasticity rules, synaptic plasticity in these models is only used to learn action probabilities. Value estimates, in contrast, are modeled as abstract variables and modified using simple reinforcement learning rules. This raises the question of whether the same plasticity models used to learn action probabilities will also function well when used to learn action values, or whether different plasticity rules are needed for each task. To address this question, we consider two simple task settings, as illustrated in Figure 1. Both settings feature both action selection and value estimation, but in each, we model only one of these components using biologically plausible plasticity mechanisms; we label them as the *action selection* and the *value estimation* settings, based on which part of the task is modeled biologically. Specifically, in the action selection setting, action probabilities are learned using explicit models of synaptic plasticity, while values are treated more abstractly; in the value estimation setting, action probabilities are treated abstractly while value estimates are learned using more explicit neural models. We then describe several models of learning based on STDP that incorporate dopaminergic modulation and evaluate their performance in both settings. We also test a variant of each setting in which the reward contingencies (i.e., which action leads to which reward) change periodically. The use of a biological model for plasticity on only one task component in each setting allows for mathematical analysis of some aspects of model behavior and for a focused evaluation of the performance of each plasticity model on specific tasks. We find that although each plasticity model does well in some cases, no model is able to succeed at both action probability learning and value learning. Thus, our results suggest that the brain, and in particular corticostriatal synapses in different regions of the striatum, may need to employ distinct, specialized plasticity mechanisms to learn different tasks.

**Fig. 1.**
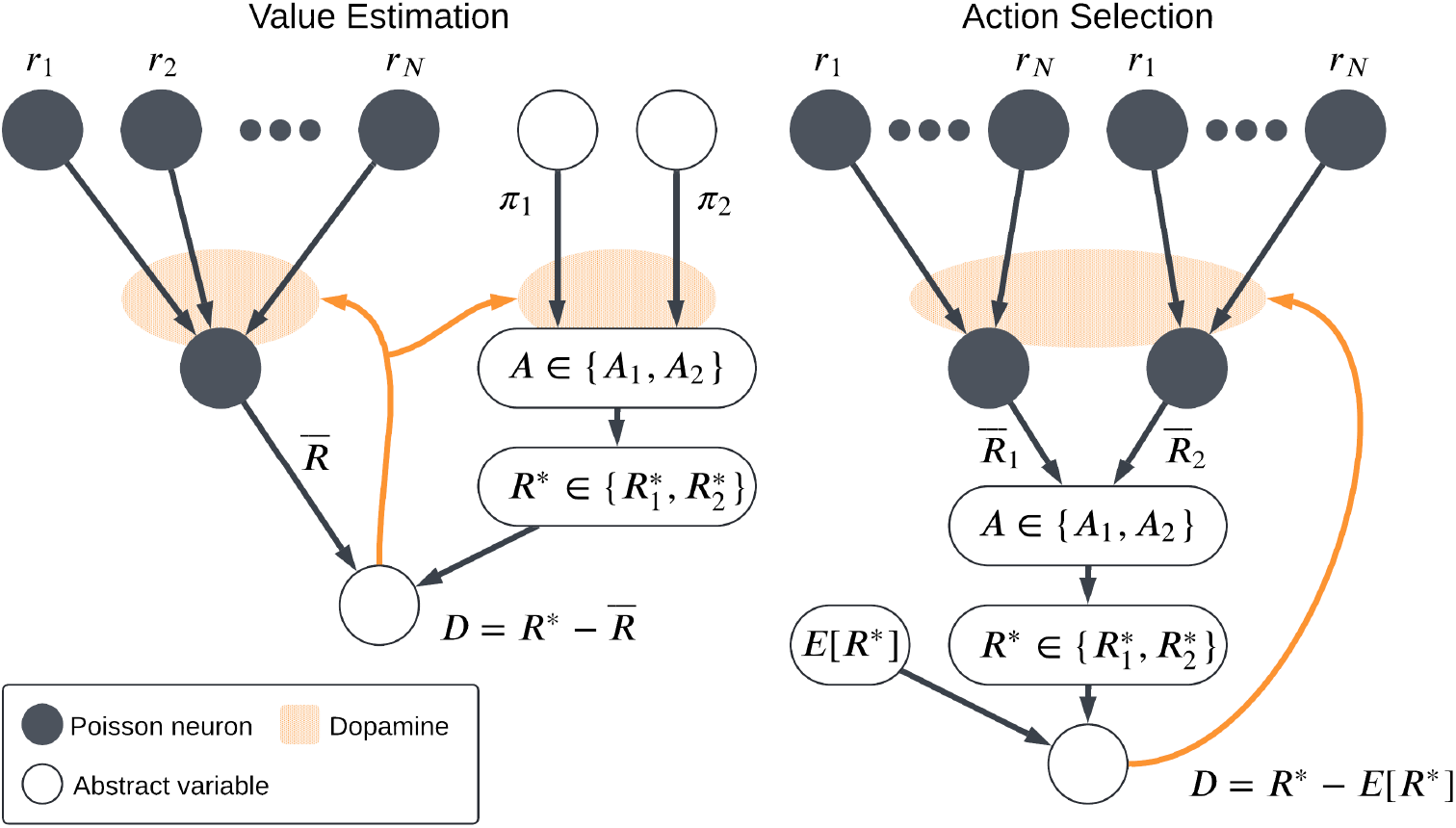
Schematic of the two main task settings. In the value estimation setting, the output firing rate 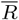 of the postsynaptic neuron is interpreted as a predicted reward and the dopamine signal *D* is the reward prediction error, while abstract variables (*π*_1_, *π*_2_) are used to determine which action is selected and hence which reward *R*\* is received. In the action selection setting, an action is chosen based on which of two competing channels has a higher output firing rate. A reward *R*\* is then received based on the action taken and the dopamine *D* again represents reward prediction error, which depends on an abstract expected reward *E*[*R*\*]

## 2 Models

### 2.1 Plasticity Models

#### 2.1.1 General Modeling Framework

Here we introduce several models of dopaminergic spike-timing-dependent plasticity. The *additive model* is based on incorporating dopamine into existing models of STDP that had been previously considered without dopamine [22, 23]. Our description and analysis of this model mainly follows the presentation in [24]. A plasticity rule known as the *multiplicative model* [25–27] was proposed to correct some deficiencies in the additive model; for reasons described below, we do not use this model, instead introducing an updated version of this plasticity rule that we call the *symmetric model*, which is better suited to modeling dopaminergic plasticity. We also study a third model, what we call the *corticostriatal model*, based on a computational model of plastic corticostriatal synaptic connections, specifically connections onto striatal spiny projection neurons that express the D1 dopamine receptor, sometimes referred to as direct pathway SPNs, as described in [21]. This model incorporates recent experimental findings about synaptic plasticity and eligibility traces in these neurons [28–34] and builds on other recent modeling studies [11–14]. To help readers to follow our exposition, we provide Table 1 as a reference for some of the mathematical notation we use.

**Table 1.** Notation for modeling framework and associated equations.

| Symbol | Meaning |
| --- | --- |
| $\rho_i^{\text{pre}}(t)$ | $i$th presynaptic spike train |
| $N$ | number of presynaptic inputs |
| $r_i, r$ | rate of $i$th presynaptic spike train, vector $(r_1, \dots, r_N)$ |
| $\rho^{\text{post}}(t)$ | postsynaptic spike train |
| $R(t)$ | rate of postsynaptic spike train |
| $\bar{R}, \bar{R}_j$ | empirical firing rate of postsynaptic neuron $j$ in some window, see (6) |
| $T_{\text{win}}$ | length of time over which $\rho_j^{\text{post}}(t)$ is averaged to compute $\bar{R}_j$ in (6) |
| $T_{\text{del}}$ | delay between time when $\bar{R}, \bar{R}_j$ are measured and when dopamine is released |
| $n_k$ | expected number of spikes by postsynaptic neuron $k$ in a window of length $T_{\text{win}}$ |
| $w_i(t), w(t)$ | synaptic weight from input $i$ to postsynaptic neuron, vector $(w_1, \dots, w_N)$ |
| $w^j$ | vector of synaptic weights $w_i^j$ to postsynaptic neuron $j$ |
| $\epsilon$ | synaptic delay |
| $\tau$ | timescale of synaptic plasticity |
| $t_{\text{pre}}, t_{\text{post}}$ | time of presynaptic or postsynaptic spikes |
| $A_i^{\text{pre}}$ | filtered signal representing spike times in $i$th presynaptic train |
| $A^{\text{post}}$ | filtered signal representing spike times in the postsynaptic train |
| $E_i^+$ | eligibility trace for modification of $w_i(t)$ based on pre-before-post spikes |
| $E_i^-$ | eligibility trace for modification of $w_i(t)$ based on post-before-pre spikes |
| $\tau_{\text{eli}}$ | timescale of eligibility trace decay |
| $D(t)$ | dopamine level, relative to a baseline |
| $r^{\text{dop}}$ | rate of dopamine release |
| $D_k, D$ | $k$th dopamine increment, released every $1/r^{\text{dop}}$ time units; index $k$ often omitted |
| $\tau_{\text{dop}}$ | timescale of dopamine decay |
| $t_{\text{dop}}$ | time when dopamine is released |
| $\lambda$ | synaptic learning rate |
| $f_{\pm}$ | model-specific synaptic plasticity functions |
| $\alpha$ | strength of negative eligibility or negative weight changes (depending on model) |
| $A_i$ | possible actions ($i \in \{1, 2\}$) |
| $p$ | probability that $A_1$ will be selected |
| $\beta$ | inverse temperature governing $p$ via equations (7) and (10) |
| $R_i^*$ | reward for selecting action $A_i$ |
| $R^*$ | actual reward received; one of $R_1^*, R_2^*$ |
| $a_{\text{sel}}$ | scaling factor for firing rate in selected channel in action selection setting |
| $\bar{A}(t)$ | 1 if previous action was $A_1$, -1 if previous action was $A_2$ |
| $\bar{\lambda}$ | timescale for changes in postsynaptic firing rate in value estimation setting, see (12) |
| $\pi_i$ | propensity to select action $A_i$ in value estimation setting; $i \in \{1, 2\}$ |

We consider linear Poisson neurons: presynaptic spike trains are modeled as Poisson processes 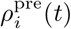 with constant rate 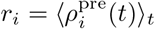 (where *i* = 1, 2, …, *N*), and likewise the spike train of each postsynaptic neuron in each model is a Poisson process 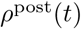 with instantaneous firing rate function *R*(*t*) given by a linear combination of the presynaptic spike trains

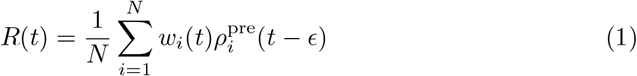

where *ϵ* > 0 is a small synaptic delay and *w*_*i*_ are the synaptic efficacies, which we will also call weights, normalized to lie in [0, 1]. We will write the vectors of input firing rates and weights as *r, w* ∈ ℝ^*N*^. This set-up can be implemented in simulations by first generating input spike trains 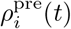, and then, whenever a presynaptic spike from input unit *i* occurs, say at time *t*_pre_, adding a postsynaptic spike at time *t*_post_ = *t*_pre_+*ϵ* to the postsynaptic spike train *ρ*^post^ with probability *w*_*i*_(*t*_pre_)*/N*. We assume that the input spike trains are uncorrelated. Note that in some cases we will have two postsynaptic neurons; then, each will have its own independent spike train, firing rate function, and so on, but since the two do not interact and hence can be treated independently, we simplify the notation by omitting subscripts for postsynaptic quantities.

In general terms, to implement synaptic plasticity, or changes in the weights, that depends on dopamine, rather than modifying each synapse immediately with the occurrence of every spike pair, as in a classical two-factor STDP rule, we instead define eligibility traces [35, 36] to track spike pairs. The eligibility traces for each synapse decay exponentially between spike pairs and are incremented when the pre- and postsynaptic neurons associated with a synapse both spike in a certain order. Weight changes are proportional to both the current values of the eligibility traces and the value of a dopamine signal, which we also model.

At a more detailed level, we base our implementation of this model on the procedure described in [14], which uses a set of trace variables to track the influences of pre- and postsynaptic spikes and spike pairs. Specifically, we define 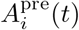 and *A*^post^(*t*) to track the pre- and postsynaptic spiking,

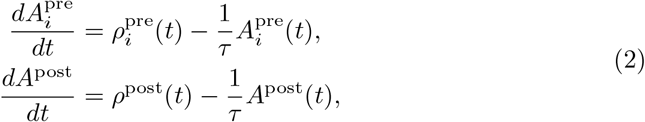

with decay constant *τ* > 0. An important assumption in our analysis, made by [24, 26] and others, is that changes in weight from individual spike pairs can be summed independently. To realize these changes, we define two eligibility traces, 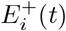 and 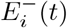, to track pre-before-post and post-before-pre spike pairs, respectively,

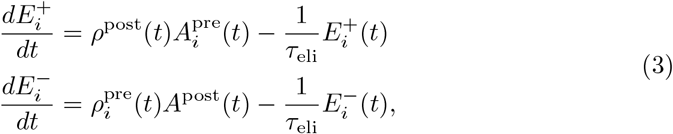

with decay constant *τ*_*eli*_ > 0. We use two independent traces in part because experimental results have suggested that this independence is present in cortical synapses [37]. Moreover, using a single trace, as done previously [14], allows spike pairs to interact nonlinearly and partially cancel each other out, while using two traces ensures that different spike pairs do not interact, which is convenient for analysis. In Appendix D we show that using a modified plasticity model with a single eligibility trace gives qualitatively similar results in most cases, and does not meaningfully improve performance on the tasks we study here.

We assume for simplicity that dopamine is released periodically at fixed intervals of length 1*/r*^dop^ for constant *r*^dop^ > 0; otherwise, it decays exponentially:

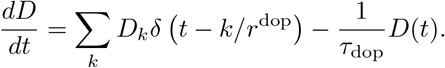

The value of the dopamine increment *D*_*k*_ depends on the task setting; see Section 2.2. We will treat this signal as the dopamine level *relative to some baseline*, rather than the raw dopamine concentration itself; so, in the absence of any signal, *D*(*t*) equals zero, and we allow *D*_*k*_ < 0. Note that while the dopamine *concentration* may depend on the postsynaptic spike train, we assume for analytical convenience that the *timing* of dopamine delivery is independent of the spiking activity. The precise form of the dopamine process is irrelevant as long as it has mean rate *r*^dop^, is independent of the spike trains, and yields dopamine signals that are far enough apart that their interactions can be neglected.

Finally, the weights in equation (1) are updated using the values of the dopamine signal and the eligibility traces in a way that depends on the choice of plasticity model. The additive, multiplicative, and symmetric models use the following rule:

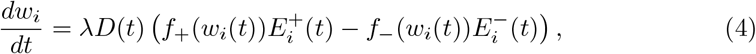

where *f*_+_ and *f*_*i*_ depend on the plasticity model used and *λ* > 0 is the learning rate. The corticostriatal model is formulated somewhat differently, as described in Section 2.1.3.

#### 2.1.2 Additive and Multiplicative Models

For the additive model, *f*_+_(*w*) = 1 and *f*_−_(*w*) = *α*, where *α* tunes how strongly negative eligibility is weighted relative to positive eligibility (typically *α* ≥ 1). This model has the advantage of simplicity; however, because *f*_±_ are independent of *w*, it allows the weights to change without bound. In past implementations of this model, the weights were artificially constrained between 0 and 1 to avoid this issue, but a model that keeps the weights bounded naturally due to its inherent dynamics would likely be a more realistic model of synaptic plasticity. This led to the development of the multiplicative model [25–27], in which *f*_+_(*w*) = 1 − *w* and *f*_−_(*w*) = *αw*. (See [24] for an exploration of models of the form *f*_+_(*w*) = (1 − *w*)^*µ*^, *f*_−_(*w*) = *αw*^*µ*^ where *µ* ∈ [0, 1] interpolates between the additive and multiplicative models.) In previous studies, which did not include the dopamine term *D*(*t*) in equation (4), the multiplicative model kept the weights bounded between 0 and 1: the 1 − *w* term shrinks positive updates from *E*^+^ near the upper boundary, while the *αw* term shrinks negative updates from *E*^−^ near the lower boundary. However, this bounding breaks down with the introduction of dopamine. If *D*(*t*) is negative, then the 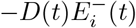 term will be positive, so the *αw* scaling factor will not prevent the weights from going above 1; similarly, the 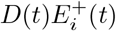 term will be negative, for which a 1 − *w* scaling factor is inappropriate for maintaining positivity of weights. The multiplicative model is therefore not effective at constraining the weights in our setting, and hence we do not consider it in our analysis.

##### 2.1.3 Alternative Models: Symmetric and Corticostriatal

One solution to this weight bounding problem is given by what we term the symmetric model, which uses *f*_+_(*w*) = *w*(1 − *w*) and *f*_−_(*w*) = *αw*(1 − *w*). (This adjustment is identical to multiplying the dynamical equations for the additive model by *w*(1 − *w*).) This choice ensures that no matter the sign of the dopamine signal or the eligibility, the weight updates will be scaled to prevent them from going past 0 or 1, as both positive and negative updates are multiplied by the same term, *w*(1 − *w*). However, whether the scaling factor *α* applies to potentiation or depression in this formulation still depends on the sign of the eligibility, and not on the dopamine.

An alternative solution, which we refer to as the corticostriatal model, is to tie the functional form of the weight update to the sign of the overall weight change, rather than just to the sign of the eligibility. In other words, we use a scaling factor of 1 − *w* when the sign of the weight change (taking into account the sign of the dopamine signal and the sign of the eligibility) is positive, and *αw* when it is negative. This convention is described by the formula

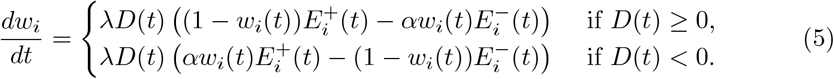

Table 2 shows how the scaling factors used by each of the four models depend on the signs of the dopamine and eligibility. The additive, multiplicative, and symmetric models only depend on the sign of the eligibility, with one factor used when *E*_*i*_ < 0 and another when *E*_*i*_ > 0, while the corticostriatal model uses the sign of the product of dopamine and eligibility to determine which scaling factor to use. Thus, in the corticostriatal model, the scaling factor corresponds to the direction in which the weights will change: 1 − *w* for increasing weights and *αw* for decreasing weights. In contrast, the scaling factors used by the other models do not correspond to the direction of weight change.

**Table 2.**
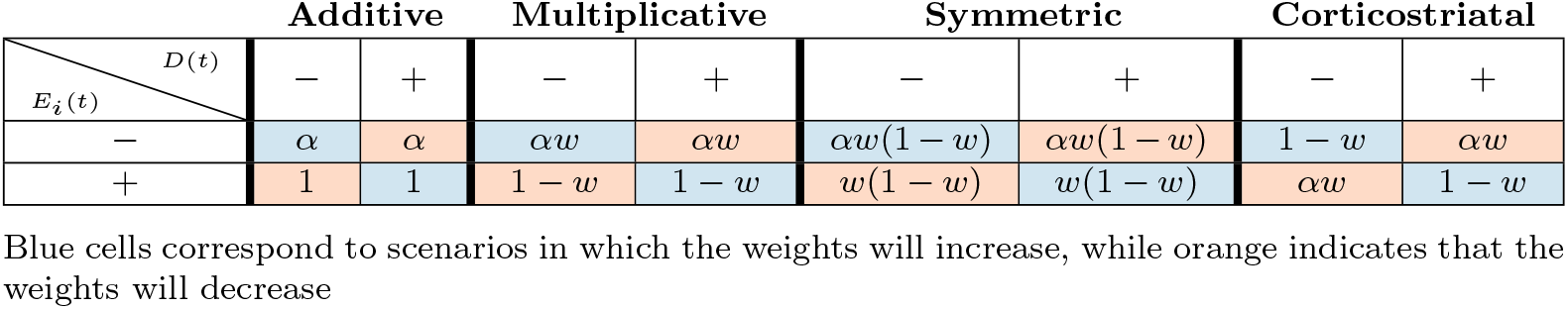
Scaling factors for the plasticity rules for positive and negative dopamine and eligibility.

To summarize, the additive model allows the weights to grow without bound (unless they are artificially constrained); the multiplicative model does not effectively solve this problem when dopamine is incorporated into the model, while the symmetric and corticostriatal models offer two different ways of effectively constraining the weights. For the remainder of this paper we focus on the additive, symmetric, and corticostriatal models.^1^ The corticostriatal model appears to be the most natural one to use in modeling corticostriatal synaptic plasticity, and it has been employed in recent modeling work [21]. As we show below, in a setting where the synaptic weights govern the likelihood that particular actions will be selected, the corticostriatal model performs well. However, it does not perform well when the weights govern value estimation. In such settings a different plasticity rule may yield an improved outcome.

Importantly, in all models, synapses become stronger with above-baseline dopamine signals (and weaker with below-baseline dopamine signals) when the postsynaptic neuron has recently participated in a pre-before-post spike pairing, and weights change in the opposite direction following post-before-pre spike pairs. These properties are implemented to match the observed behaviors of cortical synapses onto striatal spiny projection neurons expressing specifically D1 dopamine receptors [11, 13, 14, 21, 29, 32]. Neurons expressing D2 receptors show the opposite behavior, but we do not consider those here.

### 2.2 Task Settings

We consider two main task settings, corresponding to two different functional roles that striatal neurons may play: *action selection* and *value estimation*. In both the action selection and value estimation settings, the dynamics of eligibility and synaptic plasticity are as described in Section 2.1, with the dopamine signals given by equations (8) and (11), respectively.

#### 2.2.1 Action Selection Setting

In the action selection setting, we explicitly model the plasticity mechanisms that govern action selection. We implement this as a competition between two action channels [20, 38–40]. Two neurons with weight vectors *w*^1^ and *w*^2^ (with entries 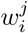 for *i* ∈ {1, 2, …, *N*} and *j* ∈ {1, 2}) receive individual, statistically independent input spike trains, with firing rates given by the same firing rate vector *r* for both *w*^1^ and *w*^2^, modeling a situation in which the two channels receive the same information from different upstream sources. A natural extension of this setting would be to employ correlated inputs, corresponding to shared presynaptic input sources, but for simplicity we do not consider this here. We compute estimates of their current firing rates 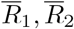 as follows:

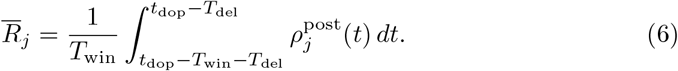

Here *T*_win_ is the length of the time window over which the spike train is averaged and *T*_del_ is a delay term between when the output firing rate is measured and the time *t*_*dop*_ when the dopamine is actually released (see Figure 2). This delay could be due to biological constraints, such as the speed of neural signal propagation or motor response, or to experimentally imposed delays.

**Fig. 2.**
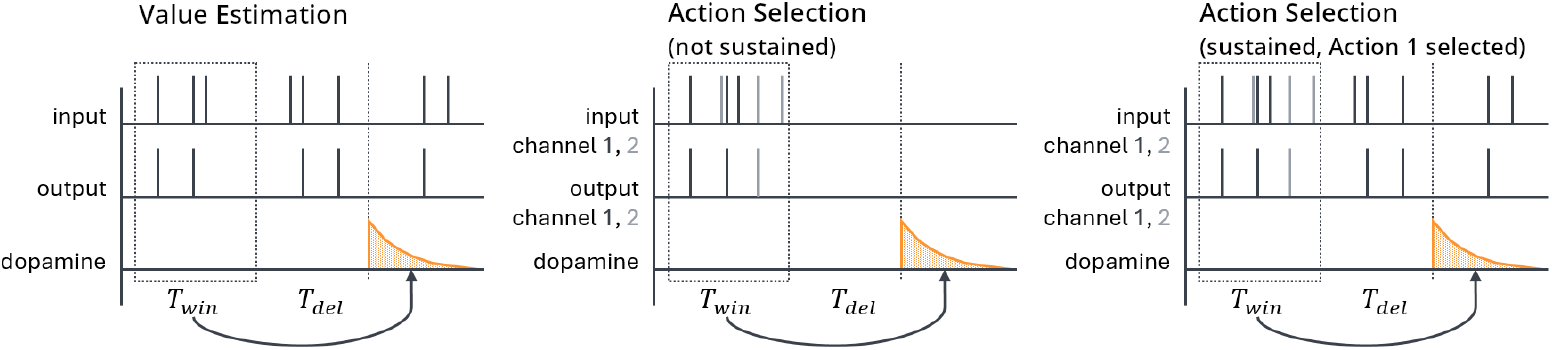
Schematic of the sequence of events in the value estimation (left) and action selection (center, right) settings. Output spikes are counted in a window of length *T*_win_ to estimate the average output firing rate; then, after a delay of length *T*_del_, dopamine is released. For the action selection setting there are two channels, colored black and gray, corresponding to the two actions being considered. In the action selection setting, we generally sustain activity in the selected channel while suppressing activity in the other channel (right), but we also test a version where activity is suppressed in both channels (center) while varying *T*_del_

The agent in this model probabilistically chooses one of two actions, *A*_1_ or *A*_2_, with selection probabilities based on the output firing rates:

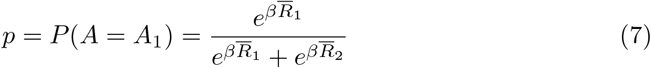

where *β* is an inverse temperature parameter and *P* (*A* = *A*_2_) = 1 − *p*. (For simplicity, in simulations in this setting we take *β* to be an arbitrary large number, so that actions are chosen deterministically based on which channel has more spikes, with ties broken randomly.) The agent receives a reward *R*\* depending on which action is taken: 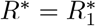 if *A*_1_ is chosen and 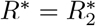 if *A*_2_ is chosen. The dopamine signal is computed as the reward prediction error, given by the difference between actual and expected rewards. In this setting, we do not model the neural mechanisms that implement value estimation and instead use an action-independent estimate of an expected reward, as also supported by some experiments [41, 42], which could be computed empirically based on experience. Thus, our dopamine equation takes the form

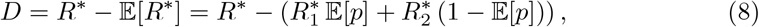

which is used to update the synaptic weights and hence 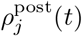 and 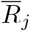, thus impacting future action selection. A successful plasticity model in this task setting is one that can learn to more frequently take the action that gives the higher reward.

In equation (8), *p* is treated as a random variable; we take the average value of *p* over instantiations of the spike trains with the given rates. That is, to compute 𝔼[*p*], we sum over the possible postsynaptic spike counts in each channel,

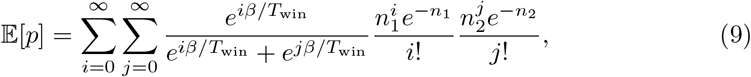

where

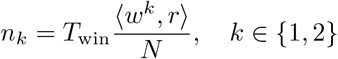

is the expected number of postsynaptic spikes in a window of length *T*_win_.

This definition assumes that the agent’s state-value function [36] is accurate. In other words, the agent has learned the reward it receives on average when performing this task with its current policy (as defined by the weights *w*^1^ and *w*^2^). The idea that value estimates are available to neurons that drive action selection is commonly used in models and has ample experimental support (e.g., [43, 44]). In practice these value estimates have to be learned, and as the animal’s policy changes, the value estimates will have to evolve along with it. In the action selection setting, we assume that the value estimates remain accurate (i.e., are learned instantaneously relative to the timescale of decision policy changes) as a simplification to allow us to focus on the action selection task without the added complication of a separate value learning circuit.

We silence all input to the striatal neuron between the end of one spike count window (used in equation (6)) and the beginning of the next (i.e., for the duration of the delay if *T*_del_ ≠ 0 as well as the period after dopamine is released before the next spike count window starts, which depends on *r*^dop^, the rate at which dopamine is released). This step is designed to represent typical experimental settings in which the input stimulus does not persist after an action is taken in response to the stimulus. For instance, in a task in which a rodent must choose which branch of a maze to follow to receive a reward, the stimulus – the sight of the junction – necessarily cannot persist after the animal has made a choice and gone down one of the branches. However, if *T*_del_ is large compared to *τ*_eli_ (equation (3)), then the eligibility will be approximately zero by the time dopamine is released, leading to minimal weight changes. We therefore generally suppose that the cortex maintains some level of activity in the channel corresponding to the selected action [45, 46] to help correctly assign credit for rewards to actions when they are separated by significant delays. Specifically, we suppose that during the interval between spike count windows, the firing rates in the selected channel equal *a*_sel_*r*_*i*_ for some *a*_sel_ ∈ (0, 1], and are zero in the channel that was not selected. We also briefly test a version of the model without this sustained activity. Figure 2 shows an illustration of the two versions of the action selection setting as well as a comparison to the value estimation setting (described below).

#### 2.2.2 Value Estimation Setting

The second task setting that we consider is what we will call the *value estimation setting*. Whereas in the action selection setting we explicitly link action selection to a comparison of postsynaptic (i.e., striatal) firing rates and leave the value estimation process fairly abstract, here we do the reverse, modeling the neural implementation of value estimation while leaving action selection abstract. We use a single postsynaptic neuron with weight vector *w*; as before, it receives independent input spike trains from a collection of presynaptic neurons with firing rates specified by entries in the vector *r*. We estimate the postsynaptic firing rate 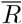 as in equation (6); in this setting, this rate represents the estimated value. To select actions, we track two variables, *π*_1_ and *π*_2_, representing how much each action is favored. These are not computed by estimating the firing rate of some neurons as in the action selection setting. They may be taken to represent the activity level of some neural population, but in the model they are simply treated as abstract variables. (Their dynamics will be described below.) We compute the probability of taking each action similarly to equation (7):

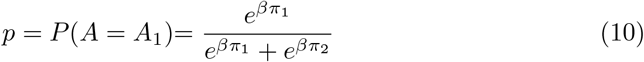

The agent receives a reward *R*\* with value 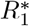 or 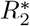 depending on whether action *A*_1_ or *A*_2_ is selected. The dopamine signal is then given by

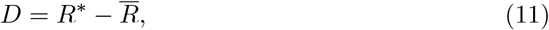

which is used to update the synaptic weights *w* using one of the plasticity rules described previously.

The dopamine signal is also used to update the values of *π*_1_ and *π*_2_. We assume that changes in these variables are proportional to the magnitude of the dopamine signal. We also assume that if action *A*_1_ is selected and the dopamine signal is positive, then *π*_1_ will increase and *π*_2_ will decrease, and vice versa if dopamine is negative or if *A*_2_ is selected. To model this, we define the variable 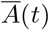 as 1 if the previous action was *A*_1_ and − 1 if the previous action was *A*_2_. Then *π*_1_ and *π*_2_ obey the following dynamical equation:

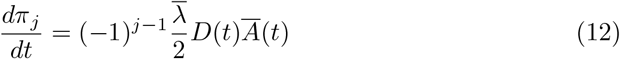

where 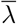 is a learning rate. (We include a factor of 1*/*2 here to eliminate a factor of 2 that arises later in our analysis.) Note that tracking both *π*_1_ and *π*_2_ is redundant, as it can be shown from equation (10) that the probability of picking each action only depends on their difference. Our simulations use a single variable, *π*_diff_ = *π*_1_ − *π*_2_, but for our mathematical analysis we find it convenient to express the system in terms of *p*, the probability of picking action *A*_1_ (see Appendix A).

Note that as there is only one channel in the value estimation setting, we do not reduce its activity level outside the spike count window using *a*_sel_ like we do in the action selection setting and instead simply keep the input firing rate constant.

### 2.3 Simulations

We use the parameter values listed in Table 3 as the defaults in our simulations for each of the two main task settings; any other parameters or changes to the defaults are listed in figure captions. “Steps” refers to the number of cycles of dopamine release in each simulation; this number as well as the learning rates *λ* and 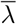 were chosen to balance noise level with computation time and to illustrate phenomena of interest. *w*_init_ = 0.5 and *p*_init_ = 0.5 were chosen because they lie at the midpoint of [0, 1] and hence represent unbiased starting values. We chose input firing rates *r* to roughly match the frequency of cortical input to the striatum. 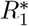 and 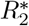 are arbitrary and were chosen for illustrative purposes. We use *α* = 1 as the default scaling parameter in our weight update equations (equations (4) and (5)) for simplicity. For the choice of *τ* = 0.02 s in equation (2), see [2] and [24]; note that [47] give values of *τ* = 0.0168 s for the decay constant of long-term potentiation (LTP) as a function of pre-before-post spike separation and *τ* = 0.0337 s as the analogous, post-before-pre quantity for long-term depression (LTD), but as we use *τ* to track the window of influence for both pre- and post-synaptic spikes, we use the intermediate value of *τ* = 0.02 s used in other sources. The half-life of dopamine has been estimated as 0.72 s in the dorsolateral striatum [48]; translating the half-life into an exponential time constant we get *τ*_dop_ ≈ 1 s. The choice of eligibility time constant *τ*_eli_ = 1 s reflects experimentally derived estimates ([33, 49]; but see [34], which finds a somewhat larger value). The delay time for dopamine release, *T*_del_, was chosen to be long enough that any spikes occurring before the delay have minimal impact on the weight changes. *r*^dop^ was likewise chosen to be small enough that the effects of any interactions between adjoining dopamine signals would be negligible. The models appear more sensitive to these parameters in the action selection setting, necessitating longer delays. The constant *β* was selected arbitrarily. In the value estimation setting we use *β* = 1; in the action selection setting we use *β* = 10^6^ to make the choice essentially deterministic for a given set of spike counts, for simplicity. The parameter *a*_sel_ was set to 0.7 to match the corresponding parameter `“sustainedfraction”` in [21].

**Table 3.** Default simulation parameters for the two main task settings.

|  | Action Selection | Value Estimation |
| --- | --- | --- |
| Samples | 100 | 100 |
| Steps | 1000 | 1000 |
| $\lambda$ | 0.01 | 0.001 |
| $\bar{\lambda}$ | N/A | 0.0025 |
| $w_{\text{init}}$ | 0.5 | 0.5 |
| $p_{\text{init}}$ | N/A | 0.5 |
| $N$ | 1 or 2 | 1 or 2 |
| $r$ | 10 or (15, 5) $\text{s}^{-1}$ | 10 or (10, 10) $\text{s}^{-1}$ |
| $R_1^*, R_2^*$ | (2, 1) | (7.5, 2.5) |
| $\alpha$ | 1 | 1 |
| $\tau$ | 0.02 s | 0.02 s |
| $\tau_{\text{dop}}$ | 1 s | 1 s |
| $\tau_{\text{eli}}$ | 1 s | 1 s |
| $T_{\text{del}}$ | 10 s | 3 s |
| $T_{\text{win}}$ | 1 s | 1 s |
| $\epsilon$ | 0.001 s | 0.001 s |
| $r^{\text{dop}}$ | $1/21 \text{ s}^{-1}$ | $1/7 \text{ s}^{-1}$ |
| $\beta$ | $10^6$ | 1 |
| $a_{\text{sel}}$ | 0.7 | N/A |

All figures use a sample size of 100 trials for numerical results besides Figure 10 and Figure 11, which use 10000; error bars and bands show standard deviations. In some phase portraits we include critical points; these were found analytically when possible, otherwise they were computed using the `scipy.optimize` library [50]. Note that some of the critical points found on the boundaries are not “true” critical points in the sense that they are not zeros of the dynamical equations. Rather, they are the result of the weights being clipped at 0 and 1. All code needed to run the simulations and reproduce the figures in the paper is available on GitHub at https://github.com/bsosis/DA-STDP.

## 3 Results

### 3.1 Action Selection Setting

#### 3.1.1 Analysis of Dynamics

We first consider a task of selecting between two actions in which the probability of picking each action is governed by one of the three plasticity rules, as described in Section 2.2. In Appendix C.1 we show that under suitable assumptions it is possible to derive a formula for the average drift of the weights over time for the additive and symmetric models in this setting:

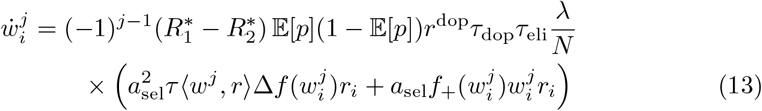

where Δ*f* = *f*_+_ − *f*_−_ and 𝔼 [*p*] is given by equation (9). An important assumption made in this derivation is that the delay *T*_del_ is large relative to *τ*_eli_; this assumption will be discussed below. We can interpret the terms in equation (13) as follows. The 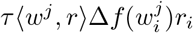 term corresponds to independent pairs of pre- and postsynaptic spikes (both pre-before-post and post-before-pre), while the 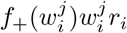 term corresponds to a pre-post spike pair in which the presynaptic spike directly causes the postsynaptic neuron to fire. Recall that when action *A*_*k*_ is selected, activity in channel *k* is reduced by a factor of *a*_sel_, while activity in the other channel is suppressed entirely. To compute 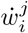 we therefore condition on action *A*_*j*_ being selected, multiplying the expression by either 𝔼 [*p*] or 1 − 𝔼 [*p*] (for actions *A*_1_ or *A*_2_, respectively), and we multiply the firing rates *r*_*i*_ in the above expression by *a*_sel_. Finally, it can be shown that the average dopamine signal, conditioned on either action *A*_1_ or *A*_2_ being selected, is 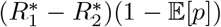 or 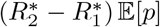, respectively. This gives the above expression.

In general, if 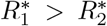 as we typically assume, then to maximally exploit the existing reward structure, we would like the difference between the weights, *w*^1^ − *w*^2^ (or ⟨*w*^1^, *r*⟩ − ⟨*w*^2^, *r*⟩ if *N* > 1), to be positive and as large as possible, so that the probability of picking action *A*_1_ is maximized. Since the weights are bounded by 0 and 1, this corresponds to the point 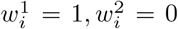 for all *i*. If 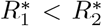 then the optimal set of weights is at 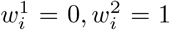. Equation (13) always has *w* = **0**, where **0** = (0, …, 0) ∈ ℝ^*N*^, as an equilibrium. For the symmetric model, it additionally has an equilibrium at *w* = **1**, where **1** = (1, …, 1) ∈ ℝ^*N*^. (When *N* > 1 the symmetric model also has a number of other equilibria, described in Appendix C.2, which we will not analyze further.) In Appendix C.2 we prove the following result about the dynamics of these models:

##### Theorem 1

*Suppose that for the additive or symmetric models in the action selection setting, the following condition is satisfied:*

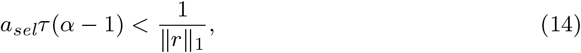

*and suppose* 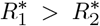. *Then the w*^1^ = *w*^2^ = **0** *equilibrium is a saddle that is repelling along the w*^1^ *coordinates and attracting along the w*^2^ *coordinates. For the symmetric model, w*^1^ = **1**, *w*^2^ = **0** *is an attractor, w*^1^ = **0**, *w*^2^ = **1** *is a repelling equilibrium, and w*^1^ = *w*^2^ = **1** *is a saddle that is attracting along the w*^1^ *coordinates and repelling along the w*^2^ *coordinates*.

These relationships are reversed if 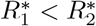. In both cases, if equation (14) holds, the flow is in the direction of the optimal set of weights for the task: 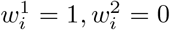 for all *i* if 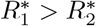 and 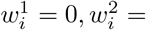 for all *i* if 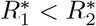.

The corticostriatal model is somewhat more complex to analyze. In Appendix C.1 we derive the following system of equations describing its average weight drift, which holds when 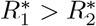:

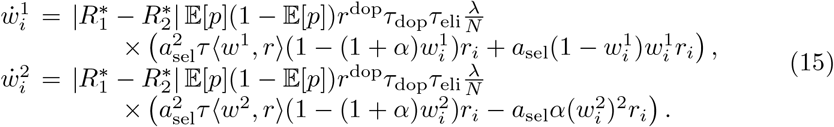

If 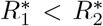, then we obtain an analogous system with the forms of the right hand sides swapped. In addition to the equilibrium point with all *w* components equal to zero, these equations have an extra equilibrium at the point (*w*^1*^, *w*^2*^) defined by

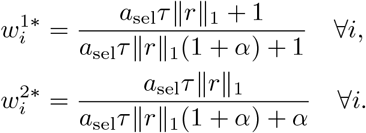

There are also equilibria at the points (**0**, *w*^2*^) and (*w*^1*^, **0**).

We show in Appendix C.2 that both *w*^1*^ and *w*^2*^ are attractors for their respective equations, so that *w*^1^ converges to *w*^1*^ and *w*^2^ converges to *w*^2*^. Since *w*^1*^ > *w*^2*^, the more rewarding action, *A*_1_, will be favored over *A*_2_, although the difference in weights after convergence may not be as great as that achievable under the additive or symmetric models.

#### 3.1.2 Simulation Results

Figure 3 shows phase portraits for the models for three different values of *α* for *N* = 1, i.e. a single weight per channel. To generate each plot, we ran a set of simulations of the fully stochastic (i.e., non-averaged) implementation (see Section 2.3) of the appropriate model with initial condition *w*_init_ = (0.5, 0.5) as marked by the × symbol. In each simulation, from this starting point, *w*^1^ and *w*^2^ evolved over 1000 time steps, and the positions of (*w*^1^, *w*^2^) at certain time steps were plotted as points with time-dependent coloring as indicated by the color bar; this process resulted in a cloud of points over many simulations, representing the distribution of weights over time. Each plot also includes all critical points, the actual trajectory from *w*_init_, and the vector field arrows for the averaged model. The orientations of these arrows indicate the directions that trajectories would move over time, from various starting positions, under the flow of the averaged model, while their lengths represent the magnitudes of the weights’ rates of change.

**Fig. 3.**
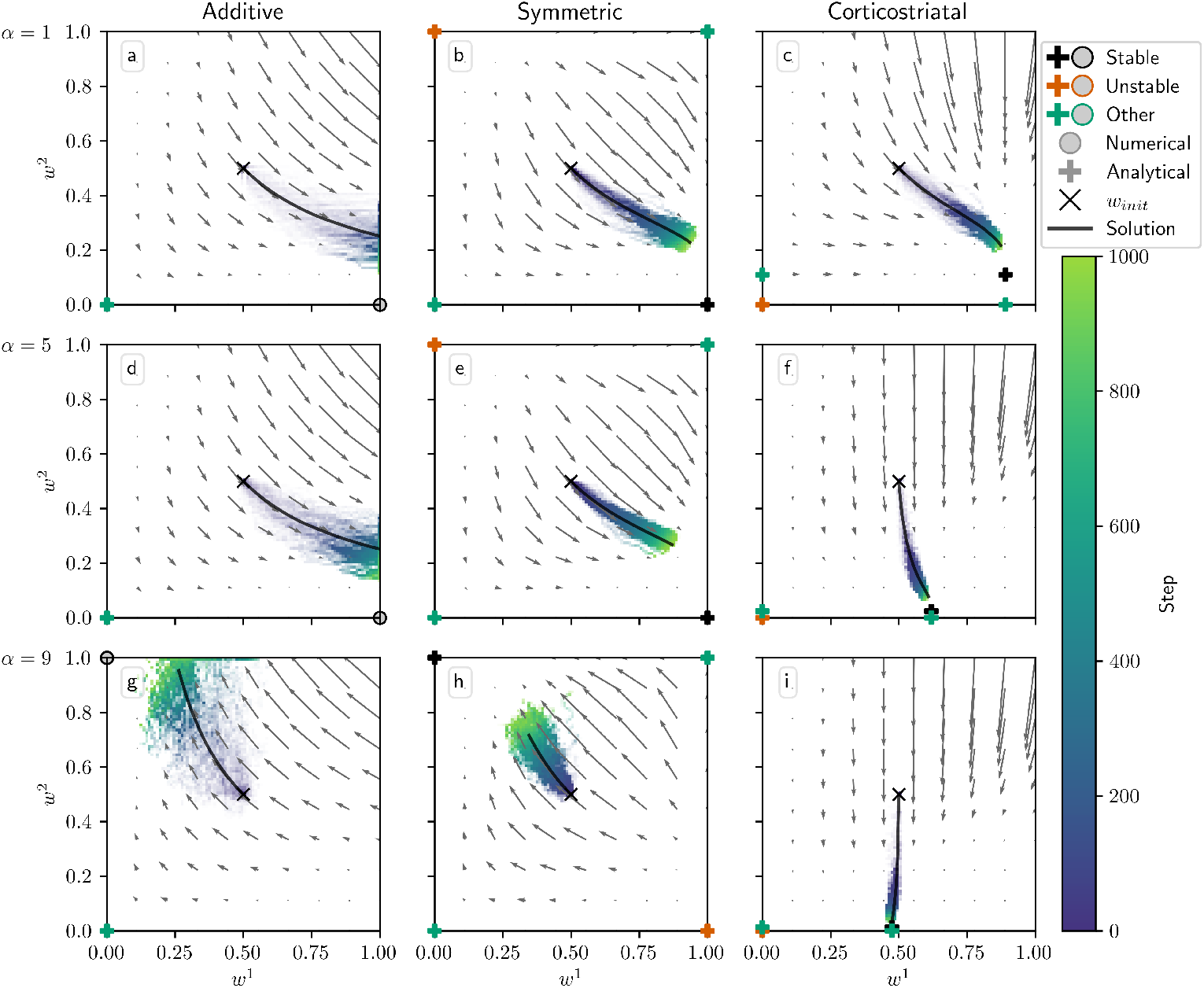
Distribution of *w*^1^ and *w*^2^ over time in the action selection setting as *α* is varied. The color code indicated in the color bar indicates the simulation step. Columns show the additive (a, d, g), symmetric (b, e, h), and corticostriatal (c, f, i) models. *α* is varied across rows: (a-c) *α* = 1; (d-f) *α* = 5; (g-i) *α* = 9. Each panel includes the model equilibria as well as arrows showing the vector field of the averaged model. Note that the vector field arrows are scaled to highlight relative flow intensities within individual plots; this scaling varies across plots. Orange coloring of the equilibria indicates unstable critical points; black indicates stable; green indicates any other equilibria. Plus markers indicate that the critical point was found analytically; circles indicate those found numerically. We use *w*_init_ = 0.5 here, marked by the “×” in each plot. The numerical solution to the averaged system starting at *w*_init_ = 0.5 is plotted in gray

We can see that the behavior of the system matches what we described above. Since 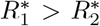, when *α* is not too large, under the additive and symmetric models, *w*^1^ increases while *w*^2^ decreases, leading to a high probability of selecting the more rewarding action, *A*_1_. However, when *α* crosses the threshold defined by equation (14), in this case *α* ≈ 8.14, the signs of the dynamical equations switch, and the system no longer selects the more-rewarded action (Figure 3g-h). Meanwhile, the weights under the corticostriatal model converge to the *w*^1*^ and *w*^2*^ equilibria defined above. Since *w*^1*^ > *w*^2*^, this leads to selection of action *A*_1_ more often than action *A*_2_ regardless of *α* (although the precise probability of selecting each action converges to a value that does depend on *α*).

Note that in the plots based on the additive model there are extra critical points at *w*^1^ = 1, *w*^2^ = 0 (for *α* = 1, 5) and *w*^1^ = 0, *w*^2^ = 1 (for *α* = 9). These are not “true” critical points in the sense that they are not zeros of the dynamical equations. Rather, they are accumulation points resulting from the clipping of the weights at 0 and 1. Such points were found numerically, and are indicated with unfilled markers (see Figure 3 legend).

In Figure 4 we plot the behavior of the system when *N* = 2. In this case there are four weights to consider: 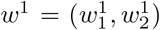 and 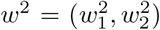. We use *r* = (15, 5) s^−1^. The optimal weight values are still at *w*^1^ = (1, 1) and *w*^2^ = (0, 0), but the asymmetry between *r*_1_ and *r*_2_ leads to different rates of convergence. We can see in Figure 4 that when *α* = 1, the behavior of the three models is qualitatively similar to that in Figure 3. When *α* = 5, though, the additive and symmetric models fail to converge to the appropriate equilibrium, because condition (14) is no longer satisfied (the condition fails at *α* ≈ 4.57 for these parameter values). Meanwhile, the corticostriatal model continues converging to *w*^1*^ and *w*^2*^, and since *w*^1*^ > *w*^2*^, this leads to selecting the correct action.

**Fig. 4.**
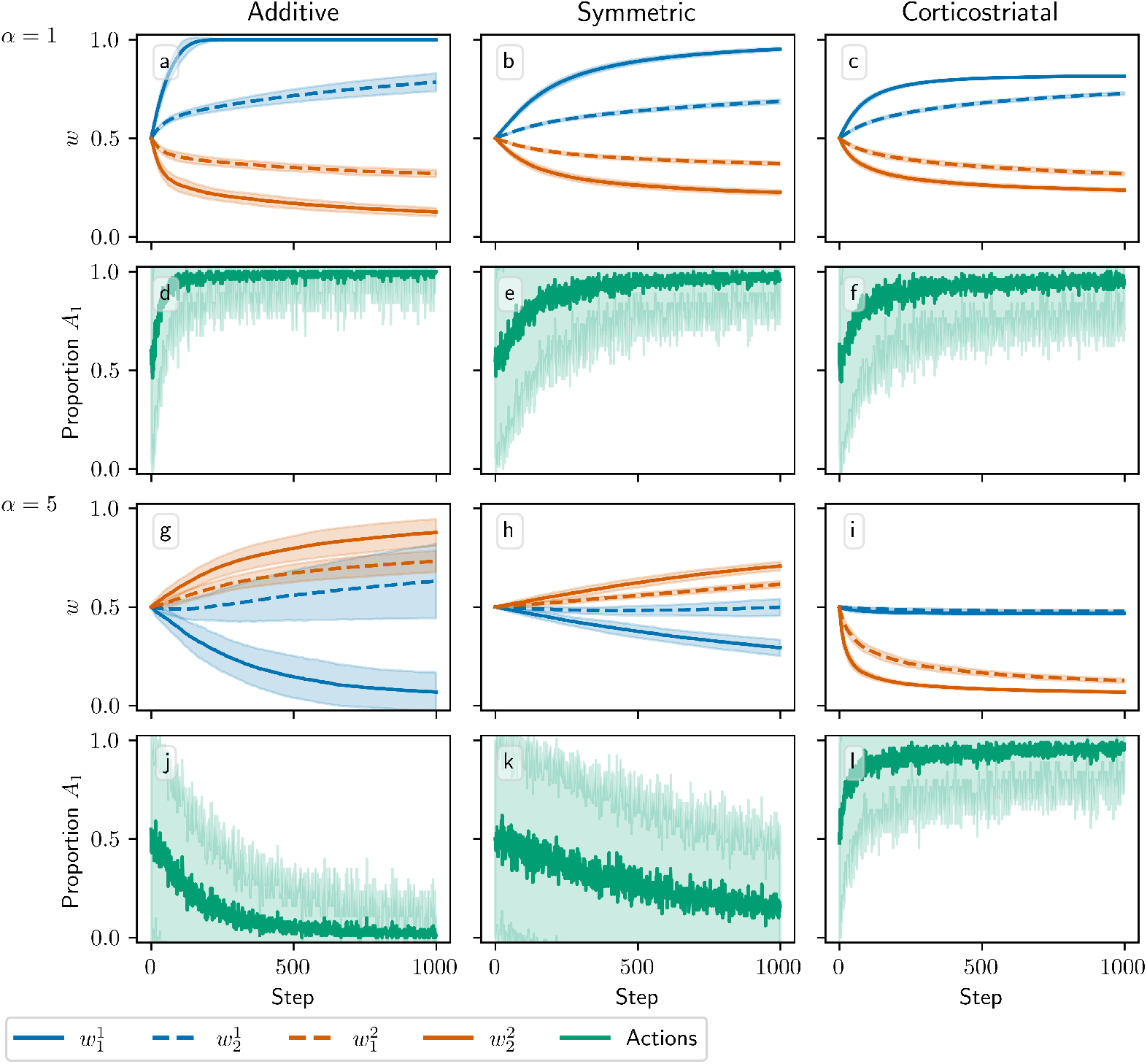
Model performance in the action selection setting as *α* is varied for *N* = 2. Plots show weights (a-c,g-i) and proportion of trials in which action *A*_1_ is selected (d-f,j-l) versus time for the additive (a, d, g, j), symmetric (b, e, h, k), and corticostriatal (c, f, i, l) models. *α* is varied across rows: (a-f) *α* = 1; (g-l) *α* = 5. Here *r* = (15, 5) s^−1^

Figures 3 and 4 show results for learning of a single relation between action and reward. In some situations, both in experiments and in natural settings, relations between actions and subsequent rewards can change over time, an effect that we refer to as contingency switching. To simulate this scenario, we swap which action is mapped to the higher reward value every 1000 steps. With this adjustment, we find that substantial differences arise in performance among the three models, as shown in Figure 5. For *α* = 1, the corticostriatal model is able to quickly react to the contingency switches and swap which action it takes, resulting in only brief drops in accuracy when switches occur. The additive model is also able to swap which action it takes, although it is significantly slower to do this and less consistent than the corticostriatal model. The symmetric model, however, is unable to perform this task well, because the weights become stuck near the widely spread values that they attain for the first contingency scenario. Increasing *α* alters the behavior of the models somewhat, but for these parameter values the corticostriatal model is able to maintain good performance.

**Fig. 5.**
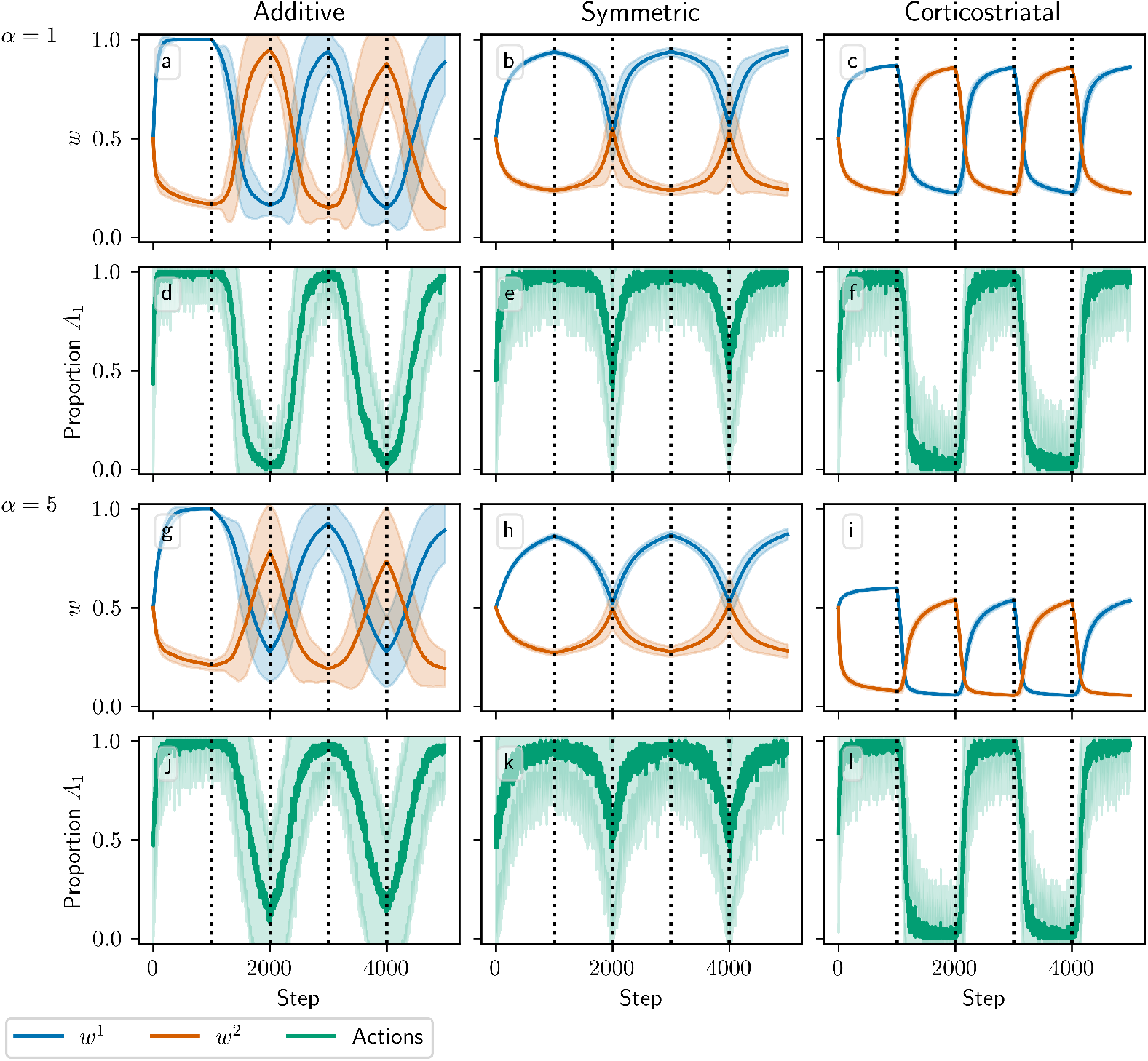
Model performance in the action selection setting with contingency switching as *α* is varied. Plots show weights (a-c, g-i) and proportion of trials in which action *A*_1_ is selected (d-f, j-l) versus time for the additive (a, d, g, j), symmetric (b, e, h, k), and corticostriatal (c, f, i, l) models. *α* is varied across rows: (a-f) *α* = 1; (g-l) *α* = 5. Here *r* = 10 s^−1^ and the reward contingencies switch between 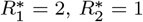 and 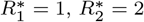 every 1000 steps

We can understand these differences in performance by referring to equations (13) and (15) and Figure 3. Swapping 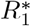 and 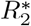 has the effect of mirroring the plots in Figure 3 across the line *w*^1^ = *w*^2^. For the corticostriatal model, this swaps between equilibria at (*w*^1*^, *w*^2*^) and (*w*^2*^, *w*^1*^). After a swap the old equilibrium is no longer a zero of the dynamical system, and because the weights are generally not too close to 0 when *α* is not too large, the drift rate will not be very small. Thus, the weights can quickly move to the new equilibrium. For the additive model, the weights tend to converge closer to 0 during the learning process. Since the weight drift equation (13) is proportional to *w*, this can cause the weights to become stuck close to the origin. The weight drift is also proportional to 𝔼 [*p*](1 − 𝔼 [*p*]), and since the action selection probability is typically higher for the additive model than for the corticostriatal model after convergence (because the difference between the weights *w*^1^ − *w*^2^ is typically greater), this term will be smaller for the additive model than for the corticostriatal model. These factors lead to slower and less consistent switching. For the symmetric model, the weight drift equation is proportional to *w*^2^(1 − *w*). This term severely slows down changes in weights near both 0 and 1, leading to very slow switching. Running the simulations with longer intervals between switches would generally not help as the weights take just as much time to escape from these values as they spend approaching them; that is, longer intervals lead to stronger convergence and hence more time needed to move away after a contingency switch.

All of our analytical results in this setting depend on a key assumption: that the delay *T*_del_ is long relative to the eligibility timescale *τ*_eli_. By imposing a large gap between when the firing rate is measured, using equation (6), and when dopamine is actually released, the delay ensures that the dopamine signal is statistically independent of the other terms in the weight update equation. This independence is what allows us to factor out the dopamine term in the average drift formulas, equations (13) and (15). Without this delay, correlations between the dopamine and the eligibility prevent us from deriving a corresponding averaged model. To judge the generalizability of our analytical results, we test model performance as *T*_del_ is varied using simulations. Figure 6 shows the weight distributions after 1000 steps for the three models as a function of *T*_del_. We also compare to an alternative task implementation without sustained activity in the selected channel. We can see that when activity is sustained in the selected channel, reducing *T*_del_ either does not meaningfully alter performance, in the case of the additive and symmetric models, or degrades performance somewhat by decreasing the difference between the weights *w*^1^ −*w*^2^, in the case of the corticostriatal model. In contrast, without sustained activity, learning is only possible when *T*_del_ is sufficiently small. In this setting, the only way for the system to learn is for eligibility from spikes within the spike count window to persist until dopamine is released. As *T*_del_ gets larger, the rate of learning decreases due to the exponential decay of the eligibility. Thus we see that using sustained activity in the selected channel effectively solves the credit assignment problem [35, 46] and results in a system that performs well when the delay is large, while without it the system is only able to successfully produce large differences between *w*_1_ and *w*_2_, and hence learn the task, when *T*_del_ is small.

**Fig. 6.**
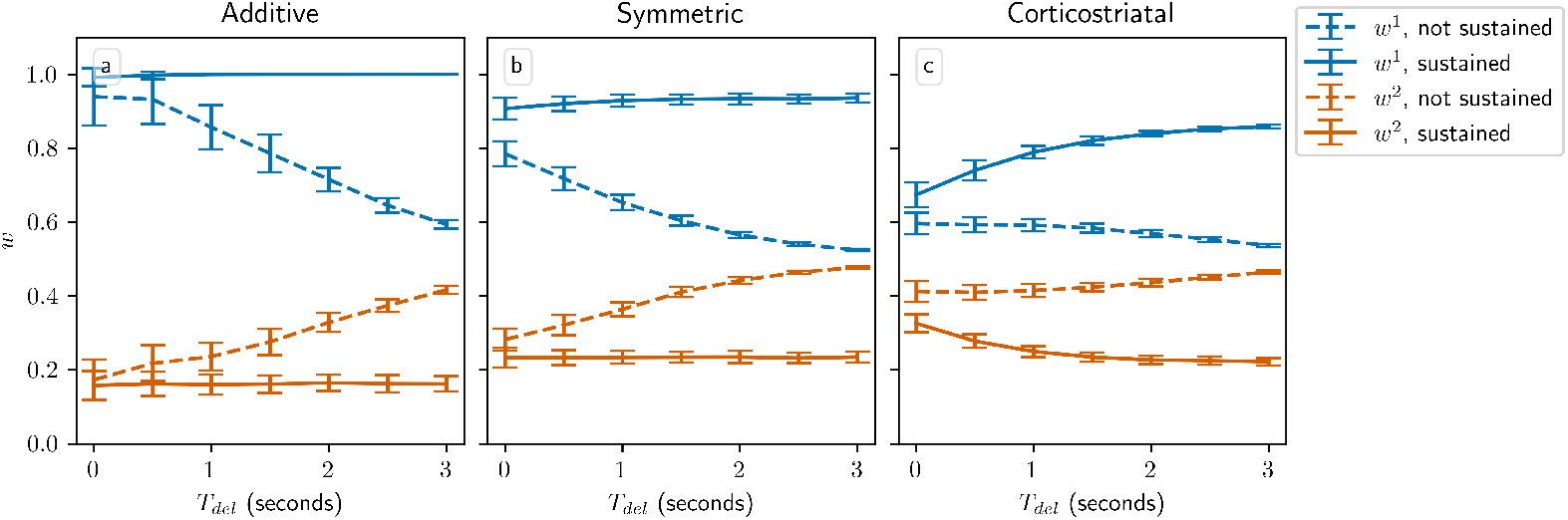
Performance in the action selection setting as delay is varied with and without sustained activity. Plots show weights after 1000 steps for additive (a), symmetric (b), and corticostriatal (c) models. With no sustained activity, both input channels are silenced during the delay period, while with sustained activity, the input to the selected channel is maintained at a level of *a*_sel_ (see Figure 2)

Overall, while the additive and symmetric models do well in the action selection setting when reward contingencies are static, the corticostriatal model does much better when they can change over time. The additive and symmetric models also seem more sensitive to *α*, with their performance tending to degrade when *α* is too large. The weakness of the corticostriatal model is that the weights tend to remain in a somewhat narrow band compared to the other two models (which can, if given enough time, drive the weights arbitrarily close to 0 and 1), leading to a lower probability of taking the correct action. This probability is determined by the number of postsynaptic spikes in each channel, and if the weights are close together, then spiking noise will sometimes lead to more spikes being counted in the incorrect channel, causing the wrong action to be taken. This outcome occurs despite the fact that here we use a large value of *β*, the temperature parameter in our action selection probability function. We believe that this problem is not a fundamental one, however, as it can be easily solved through downstream integration over the outputs of multiple striatal neurons to obtain a clearer signal. Consequently, we conclude that overall, the corticostriatal model is better suited to the action selection setting than are the additive and symmetric models.

### 3.2 Value Estimation Setting

#### 3.2.1 Analysis of Dynamics

The strong performance of the corticostriatal model in the action selection setting with contingency switches highlights its flexibility and raises the question of whether it is equally suited to learn to estimate choice values, or whether other plasticity rules will perform better for such tasks. We attempt to answer this question using the value estimation task described in Section 2.2. Under suitable assumptions it is possible to derive a formula for the average drift of the weights and action probabilities over time for the additive and symmetric models in this setting:

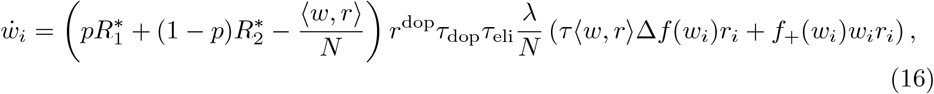

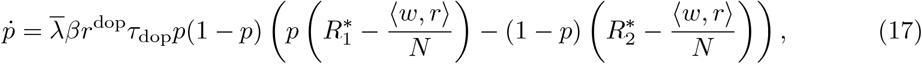

where Δ*f* = *f*_+_ − *f*_−_ and *p* is the probability of selecting action *A*_1_. See Appendix A for the derivation of system (16,17). As in the action selection setting, we assume that the delay *T*_del_ is large relative to *τ*_eli_.

The terms in equations (16) and (17) have simple interpretations. In equation (16), the *τ* ⟨*w, r*⟩ Δ*f* (*w*_*i*_)*r*_*i*_ and *f*_+_(*w*_*i*_)*w*_*i*_*r*_*i*_ terms are identical to those in the action selection setting and correspond to independent spike pairs and pre-post pairs where the presynaptic spike causes the postsynaptic spike, respectively. The 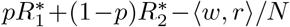 term is the average dopamine level, which is the difference between the average reward received and the average output firing rate. In equation (17), *p*(1 − *p*) arises from the logistic dependence on *π*_1_ − *π*_2_ in equation (10), and the other term describes the expected difference in dopamine levels between trials where action *A*_1_ is selected and those where *A*_2_ is selected.

System (16,17) has a number of equilibria. The following are critical points for every choice of *f*_+_, *f*_−_:

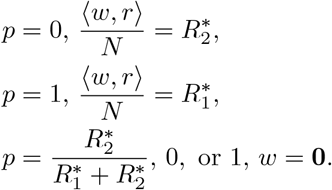

Note that the first two families of equilibria comprise (*N* − 1)-dimensional hyperplanes of weights at fixed *p*. These correspond to sets of weights such that the average output firing rate ⟨*w, r*⟩*/N* equals the average reward received. (In general, for other values of *p*, the set of weights where 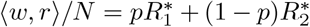 will have zero weight drift, 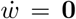, but 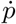 may not be zero.) Different choices of *f*_+_, *f*_−_ may lead to additional equilibria. In particular, the symmetric model has a set of extra critical points at:

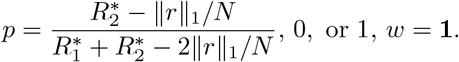

It also has a number of other equilibria which appear when *N* > 1, which we describe in Appendix B.3.

Assuming 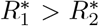, so that action *A*_1_ is more rewarding than *A*_2_, we would like *w* and *p* to converge to the *p* = 1, 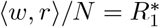 set of equilibria, at which the agent always picks *A*_1_ and the system accurately predicts the reward. Conversely, if 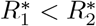 we would like the system to converge to the *p* = 0, 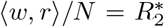 set of equilibria. In Appendix B.1 we prove the following result providing a condition relevant to the stability of these states:

##### Theorem 2

*Pick w and p so that either p* = 1, 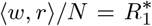 *or p* = 0, 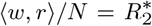. *For the additive and symmetric models given by equations* (A9) *and* (17), *the Jacobian at this equilibrium has one nonzero eigenvalue, which has an eigenvector that is zero in the p coordinate. This eigenvalue is negative if and only if*

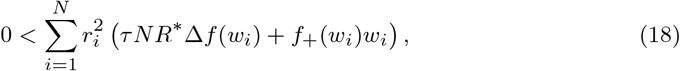

*with the choice of* 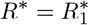 *or* 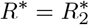 *corresponding to the equilibrium under examination*.

At these equilibria there also exists a center manifold with nonzero component in the *p* coordinate. The union of these manifolds over the equilbrium hyperplanes forms an *N*-dimensional surface, which in many cases appears to connect the two sets of equilibria at *p* = 0 and *p* = 1 (see Figure 7). We show in Appendix B.1 that the flow along this manifold depends on the sign of 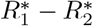:

**Fig. 7.**
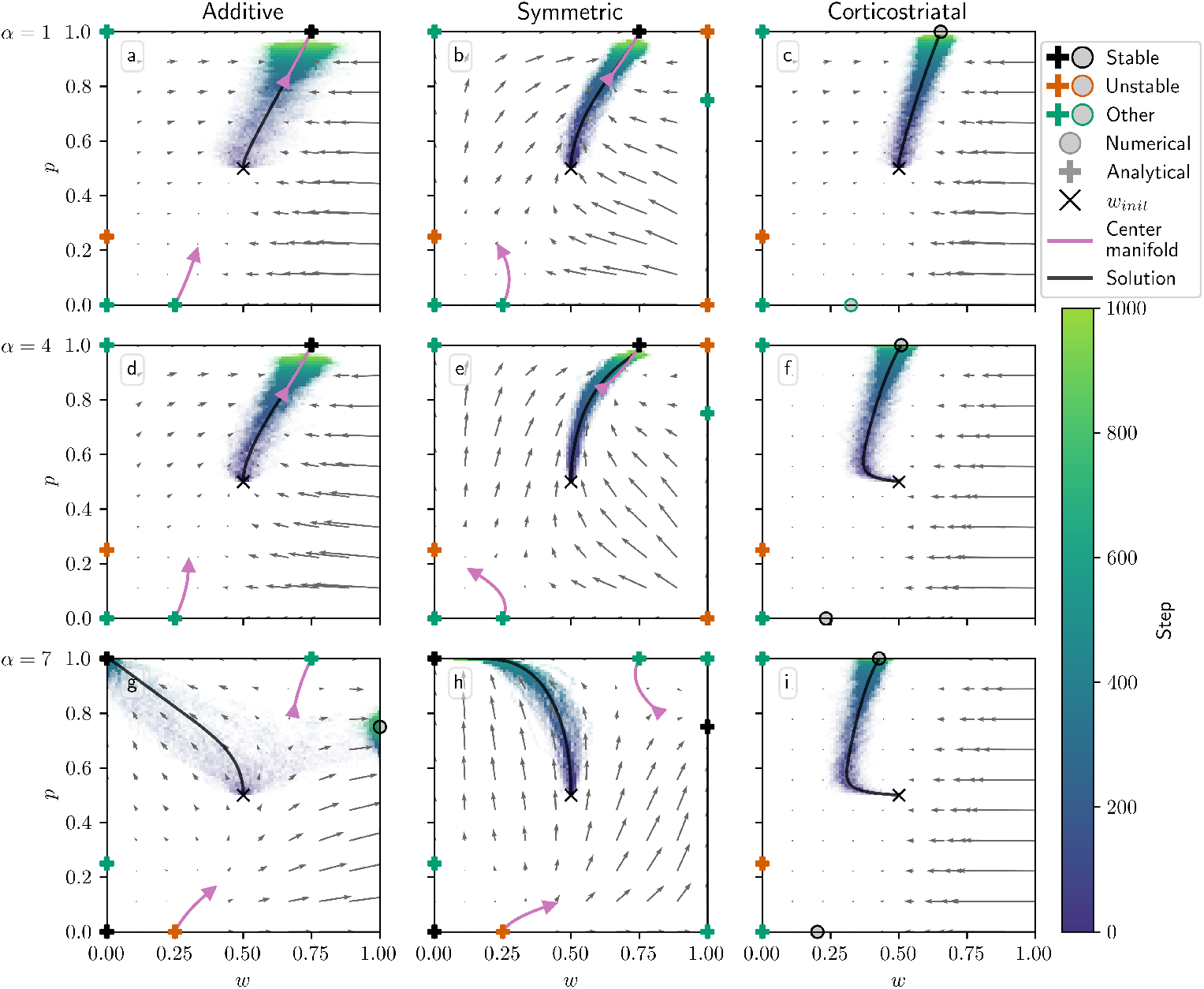
Distribution of *w* and *p* over time in the value estimation setting as *α* is varied. Columns show the additive (a, d, g), symmetric (b, e, h), and corticostriatal (c, f, i) models. *α* is varied across rows: (a-c) *α* = 1; (d-f) *α* = 4; (g-i) *α* = 7. We use *w*_init_ = 0.5, *p*_init_ = 0.5 here, marked by the “×” in each plot. The low-order shape of the center manifolds for the additive and symmetric models at certain critical points is plotted in pink. The numerical solution to the averaged system starting at *w*_init_ = 0.5, *p*_init_ = 0.5 is plotted in gray and closely aligns with the center manifold in (a,b,d,e)

##### Theorem 3

*Points of the form p* = 0, 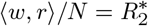 *or p* = 1, 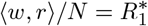 *are equilibria of equations* (A9) *and* (17) *with a nontrivial center manifold; the flow along this manifold is towards p* = 1 *and away from p* = 0 *if* 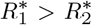, *or away from p* = 1 *and towards p* = 0 *if* 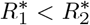.

Thus, as long as equation (18) is satisfied for a sufficient proportion of the hyperplane, the system will generally move along the center manifold towards the appropriate set of equilibria, where the more rewarding action is always picked and the reward is predicted accurately. The presence of other equilibria may complicate this picture in some cases, though. In Table 4 we summarize the equilibria of the additive and symmetric models and the conditions determining their stability; see Appendix B for proofs of all of these results. When *N* > 1, the symmetric model has a number of additional equilibria not included in the table that we do not analyze in detail; see Appendix B.3.

**Table 4.**
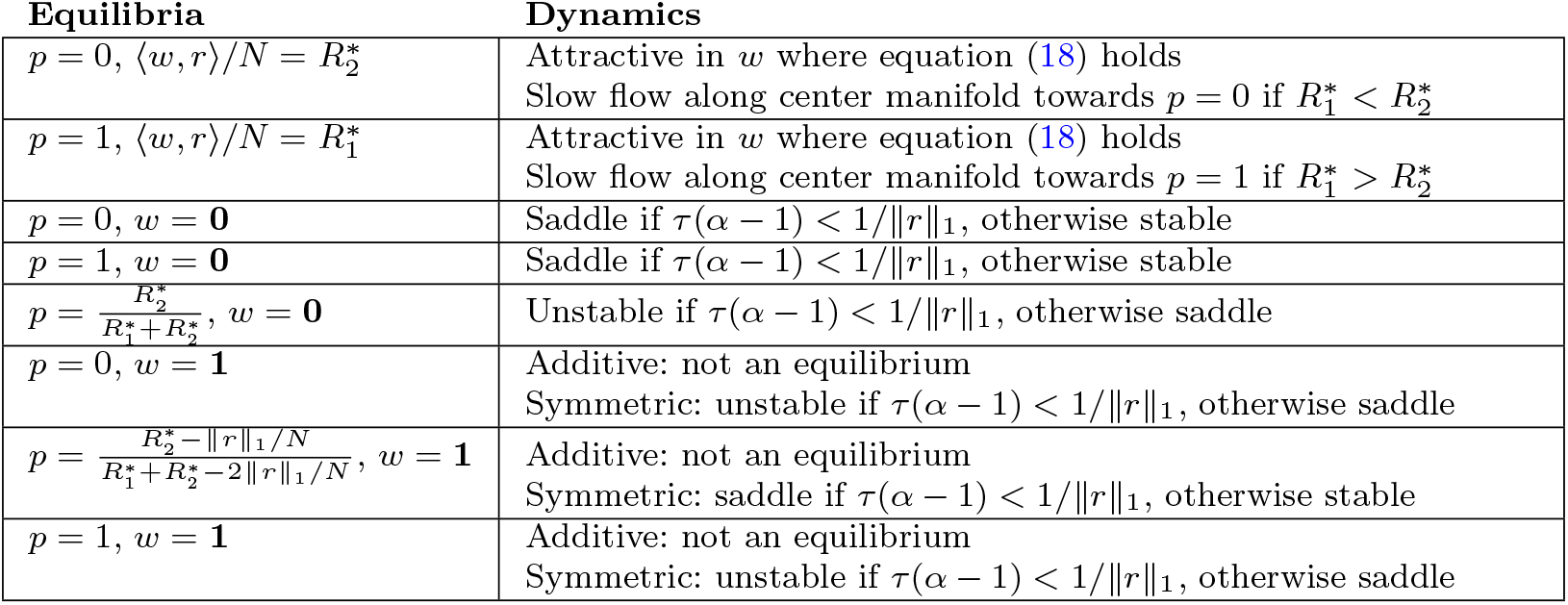
Equilibria of the additive and symmetric models in the value estimation setting.

The corticostriatal model is more difficult to analyze. Because the form of the plasticity rule in that model depends on the sign of the dopamine signal, in general it is not possible to factor out the average dopamine level 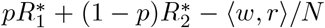 like we can for the additive and symmetric models. We analyze this model further in Appendix A.2 and Appendix B.4; its equation for the dynamics of *p* is identical to equation (17), but that for *w* differs from equation (16). In general, points on the hyperplanes 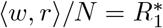 or 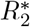 will not be critical points under this model, meaning that the weights will generally not converge to the optimal values.

#### 3.2.2 Simulation Results

Figure 7 illustrates the evolution of (*w, p*) with *N* = 1 for all three plasticity models, overlaid on the phase portraits for the corresponding averaged models, for three different values of *α*. Here we plot *w* along the *x*-axis and *p* along the *y*-axis. In addition to the averaged model vector fields, we also include a representation of the lower-order behavior of the center manifolds described above.

We can see several of the changes in behavior described in Table 4 as *α* increases. For the additive model, when *α* is not too large, *p* converges to 1 and *w* converges to 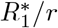, so that the predicted reward equals 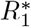. (Recall that 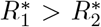, so that action *A*_1_ is more rewarding.) However, for large enough *α* (in this case *α* ≥6) the condition given in equation (18) fails, and the equilibrium loses stability. For the additive model in the *N* = 1 case this condition is equivalent to *τ* (*α* − 1) < 1*/*∥ *r*∥ _1_, so as described in Table 4, the three equilibria along *w* = 0 also change their stability as *α* increases. For the initial conditions and parameters we used, this leads to most trajectories converging to *p* = 1, *w* = 0 for *α* = 7.

Note that an extra critical point arises in the *α* = 7 plot at *p* ≈ 0.75, *w* = 1. Like we saw in the action selection setting, this is not a “true” critical point but rather the result of the weights being clipped at 0 and 1. In this case, it can be seen by examining equation (17) that at *w* = 1 and 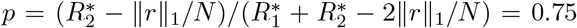, using the parameters listed in Table 3, we have 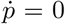 and 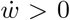, so preventing the weights from increasing above 1 makes trajectories accumulate at this point.

The symmetric model behaves very similarly to the additive model, except that it has three extra equilibria at *w* = 1. For low values of *α*, these are unstable or saddles, and the flow is away from *w* = 1. Past *α* = 6 when the *τ* (*α* − 1) < 1*/* ∥*r*∥ _1_ condition fails, *w* = 1 becomes attractive, and in particular the fixed point at 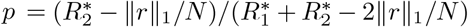, *w* = 1 becomes stable. While in the simulation shown in Figure 7h trajectories generally do not converge to it, the vector field indicates that under different initial conditions trajectories will converge to it rather than to the *p* = 1, *w* = 0 equilibrium.

For the corticostriatal model, while we are able to analytically determine the stability of the critical points along the *w* = 0 axis (see Appendix B.4), we are unable to find a closed-form expression for the two critical points evident in each of Figure 7c,f,i for nonzero values of *w*; these are instead computed numerically. Note that these *are* zeros of the underlying dynamical equations, unlike the extra equilibria we see in the additive model; they simply cannot be expressed analytically. Both the upper and lower critical points appear to remain stable as *α* increases, although their position shifts with *α*. In general, they will not equal 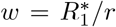 or 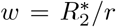, so while the system may learn to pick the more rewarding action (based on *p* → 1), the value estimates made by the system will be incorrect.

For the additive and symmetric models, we also plotted curves to indicate the low-order shape of the center manifolds at the *p* = 0, 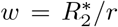 and *p* = 1, 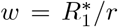 critical points (Figure 7, purple curves). As can be seen from comparison with the vector field, especially in the symmetric model, the curves describe the local behavior of trajectories near these equilibria. Note that the lengths of these curves were selected for visualization of the manifolds’ shapes and should not be taken to indicate the regions where this low-order approximation holds; for example, for *α* = 7, the vector field and center manifold align only on a very small neighborhood of the *p* = 1 equilibria.

In Figure 8, we plot the dynamics when *N* = 2, starting from three different initial conditions to better illustrate the dynamics. In this case the sets of equilibria 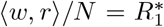 and 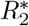 form lines in the *w*_1_ − *w*_2_ plane, indicated in the figure with dashed lines. We can see that for the additive and symmetric models, the weights converge to the 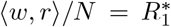 line as *p* converges to 1, indicating that the more rewarding action is being picked more often, and that the system correctly estimates the values of its rewards. There does appear to be some diffusion along this line, especially for the additive model. For the corticostriatal model, while *p* converges to 1, the weights appear to converge to a point somewhat off of the line; as we saw in the *N* = 1 case, the value estimates made by the system will be incorrect.

**Fig. 8.**
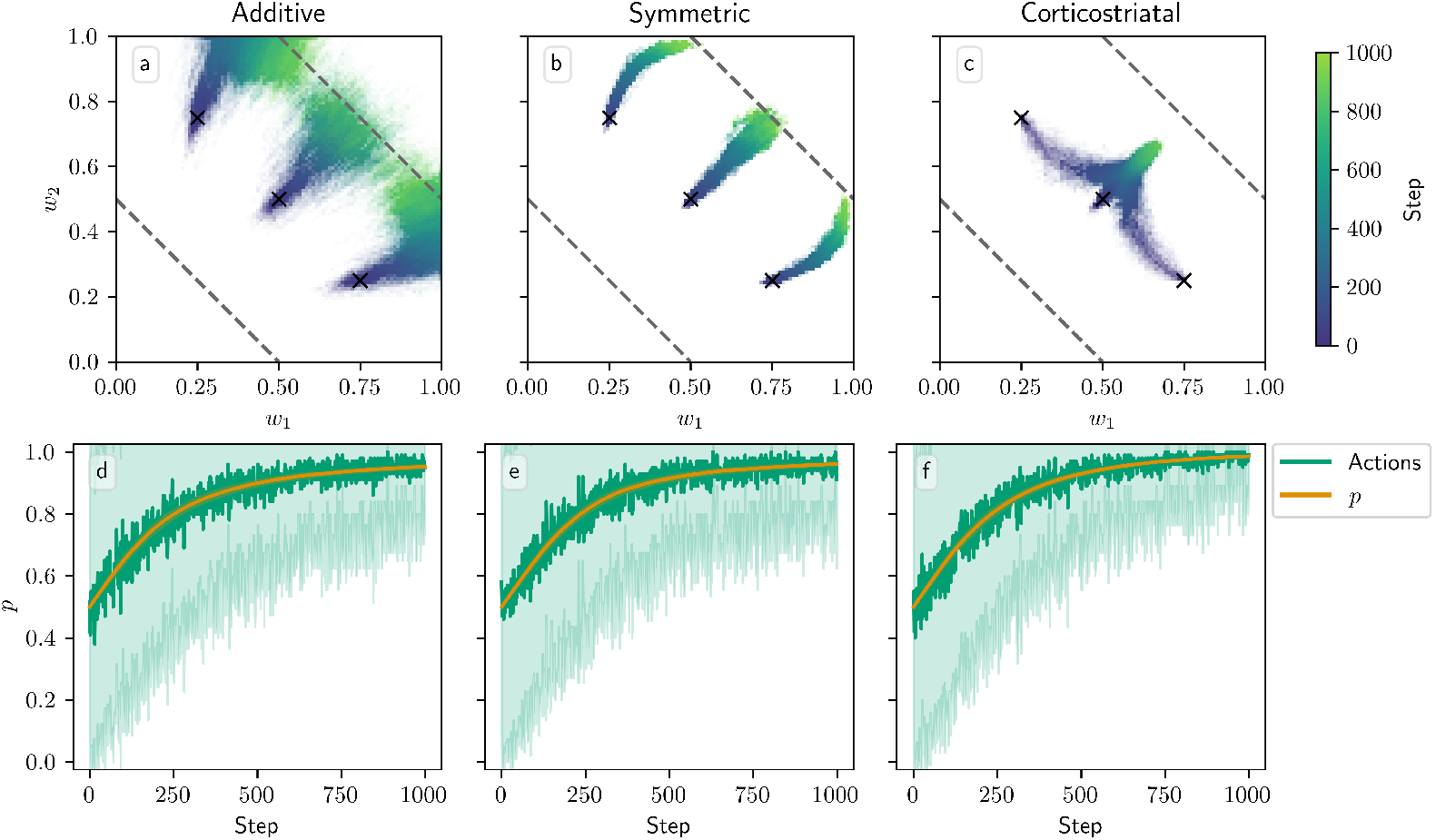
Distribution of *w* and *p* over time in the value estimation setting with *N* = 2. The color code indicated in the color bar shows the simulation step. Columns show the additive (a, d), symmetric (b, e), and corticostriatal (c, f) models. The top row shows the two weights, *w*_1_ and *w*_2_, while the bottom row shows the proportion of trials on which action *A*_1_ is picked along with the values of *p*. The two sets of equilibria at 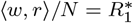 and 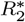 are indicated with dashed lines. (Note that these are not equilibria for the corticostriatal model; we plot the lines there only for ease of comparison.) We use *α* = 1 and three different initial conditions: *w*_init_ = (0.25, 0.75), *w*_init_ = (0.5, 0.5), and *w*_init_ = (0.75, 0.25) (all with *p*_init_ = 0.5), indicated by the “×” marks in the upper plots; the lower plots only include *p* values and action proportions for *w*_init_ = (0.5, 0.5) as the other initial conditions give almost identical results. In the lower plots, the shaded envelopes show standard deviations while solid lines show means over 100 trials

In Figure 9 we test a version of the value estimation task with contingency switching, in which we switch which action leads to which reward every 1000 steps. In general, performance with contingency switching depends on the balance between the learning rates for *w* and *p, λ* and 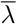. The learning rates need to be large enough that both *w* and *p* have enough time between contingency switches to approach close to their equilibrium values, but ideally not so large that the system is too noisy. Additionally, performance is significantly degraded when *λ* is large relative to 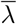. For instance, suppose 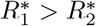, and we switch from a state in which action *A*_1_ leads to reward 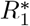 and *A*_2_ leads to 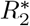 to one where *A*_1_ leads to 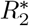 and *A*_2_ leads to 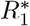. In the original state, *A*_1_ is more highly rewarded, so suppose *p* ≈ 1 and 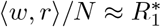, matching the convergence behavior of the additive and symmetric models. If *λ* is too large relative to 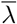, then after the switch, *w* may rapidly converge to 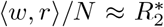 while *p* remains close to 1. Since the system is now accurately predicting the low reward it is receiving, the dopamine signal will be small, and so *p* will take a long time to leave the vicinity of 1. For this reason the system generally performs better with contingency switching when 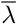 is large relative to *λ*.

**Fig. 9.**
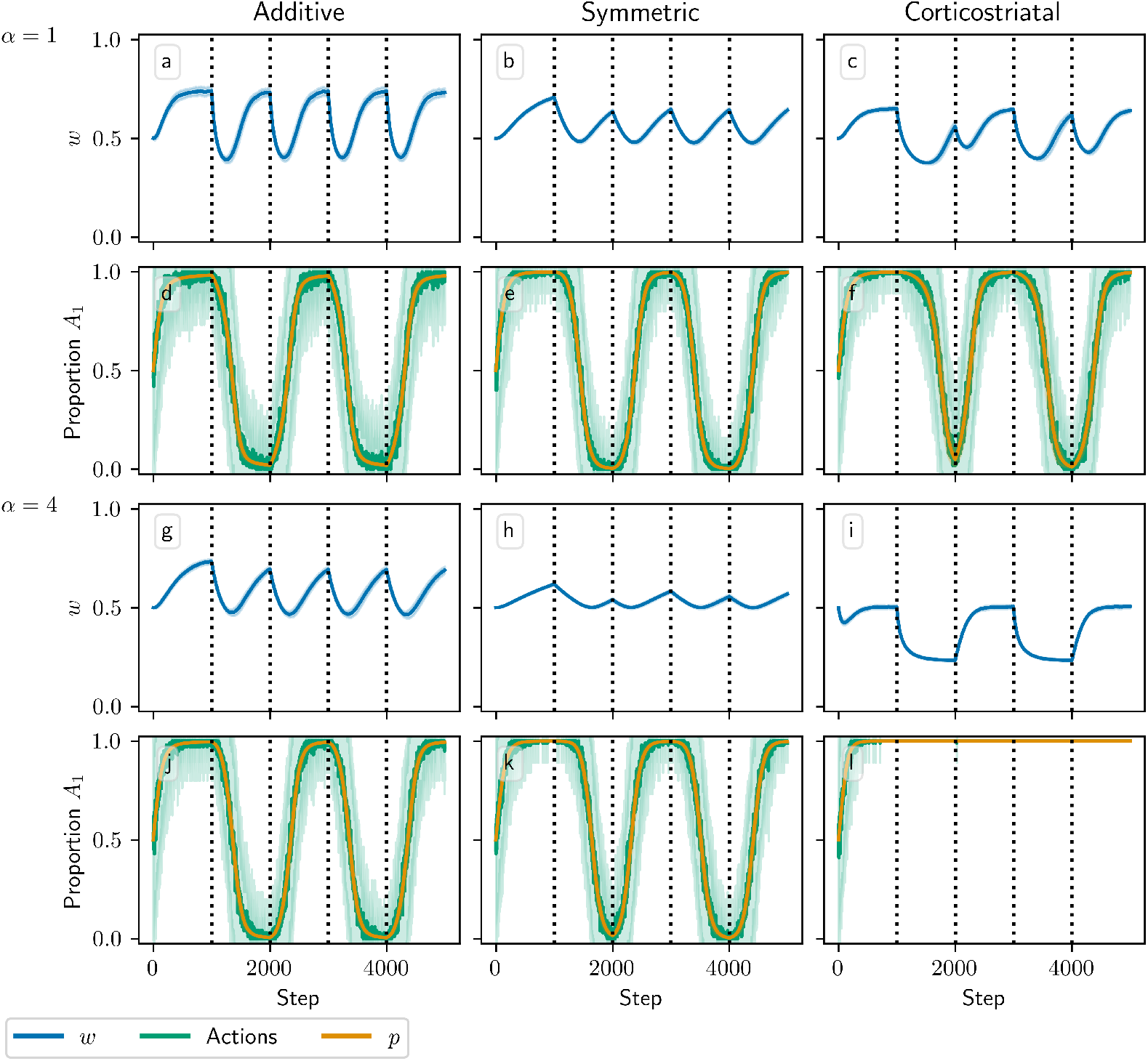
Model performance in the value estimation setting with contingency switching as *α* is varied. Plots show weights (a-c, g-i) and the proportion of trials on which action *A*_1_ is picked along with the values of *p* (d-f, j-l) versus time for the additive (a, d, g, j), symmetric (b, e, h, k), and corticostriatal (c, f, i, l) models. *α* is varied across rows: (a-f) *α* = 1; (g-l) *α* = 4. Here reward contingencies switch between 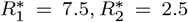 and 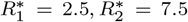 every 1000 steps; we use *λ* = 0.00015 and 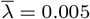 for illustrative purposes

Another potential issue occurs if the weights do not approach 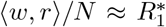 or 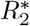 closely enough between contingency switches. This pitfall always arises with the corticostriatal model, as these hyperplanes are not equilibria for this model, but it also may occur in other cases if *λ* is too small. If the weights do not converge properly between switches, then *p* may converge to 0 or 1 *too strongly*. To see this, observe that near *p* = 0 or *p* = 1, 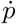 is proportional to the estimated value term 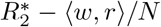 or 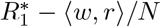, respectively (see equation (17)). If *w* is close to its equilibrium value, then this term will be small, and generally *p* will not approach 0 or 1 too closely. However, if it does not converge properly, the estimated value term will remain large and *p* will be driven to 0 or 1 exponentially, as 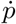 will be proportional to *p*(1 − *p*) times a factor significantly different from zero. This is problematic in contingency switching tasks because it may then take too long for *p* to leave this point when contingencies swap.

Due to these considerations, each of the three plasticity rules needs different values of *λ* and 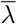 to perform well. In Figure 9 we use *λ* = 0.00015 and 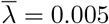, in which all three models do at least moderately well at selecting the correct action when *α* = 1, although the symmetric and corticostriatal models do not do as well at accurately predicting their rewards. A higher *λ* would help the symmetric model but harm the corticostriatal model, which would benefit from a lower *λ*, and all three models require a higher 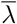 than the value of 0.0025 used in other figures. Performance of these models also degrades significantly for larger values of *α*, although different choices for *λ* and 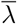 may ameliorate this.

All of our analytical results in this setting depend on the assumption that the delay *T*_del_ is long relative to *τ*_eli_, like in the action selection setting. This assumption is what allows us to factor out the dopamine term 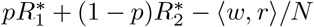 in the average drift formula, equation (16). Without this term, we can no longer guarantee that points on the hyperplanes 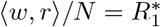 or 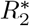 are equilibria for any of the three plasticity models. In Figure 10 we plot the change in weight after a single dopamine release as a function of *w*_init_ in an *N* = 1 setting, with *p*_init_ = 0.5 fixed, while varying *T*_del_. When *T*_del_ = 3 s the simulations obey the predictions of the averaged models, and in the additive and symmetric cases they intersect the x-axis at *w* ≈ 0.5, the point at which 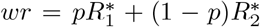 when *p* ≈ 0.5. (While the plots may appear fairly noisy, keep in mind that they only display the change in weight after a single dopamine signal. Figure 7 shows that although there is some dispersion, over the course of many trials trajectories still tend to follow the averaged dynamics.) When *T*_del_ = 0 s the simulations do not exactly match the predictions, but the differences are fairly small; they are most visible for the symmetric model. In most cases where they differ the *T*_del_ = 0 s curves are below the curves for *T*_del_ = 3 s. This undershoot may occur because there is a source of negative correlation between the dopamine value and the eligibility at the time that dopamine is released. Specifically, the dopamine value *D* from equation (11) is negatively correlated with the number of postsynaptic spikes in the spike count window, while the eligibility will in most cases (depending on the plasticity model and the parameters) be positively correlated with the number of recent spikes. If there is no delay, then this will include the spikes in the spike count window used to compute the dopamine value. While these plots show that for realistic parameter values our model is not very sensitive to the delay or its absence, it should be noted that for other sets of parameters, for instance smaller values of *τ*_dop_, *τ*_eli_, and *T*_win_, the lack of a delay can have a significant effect.

**Fig. 10.**
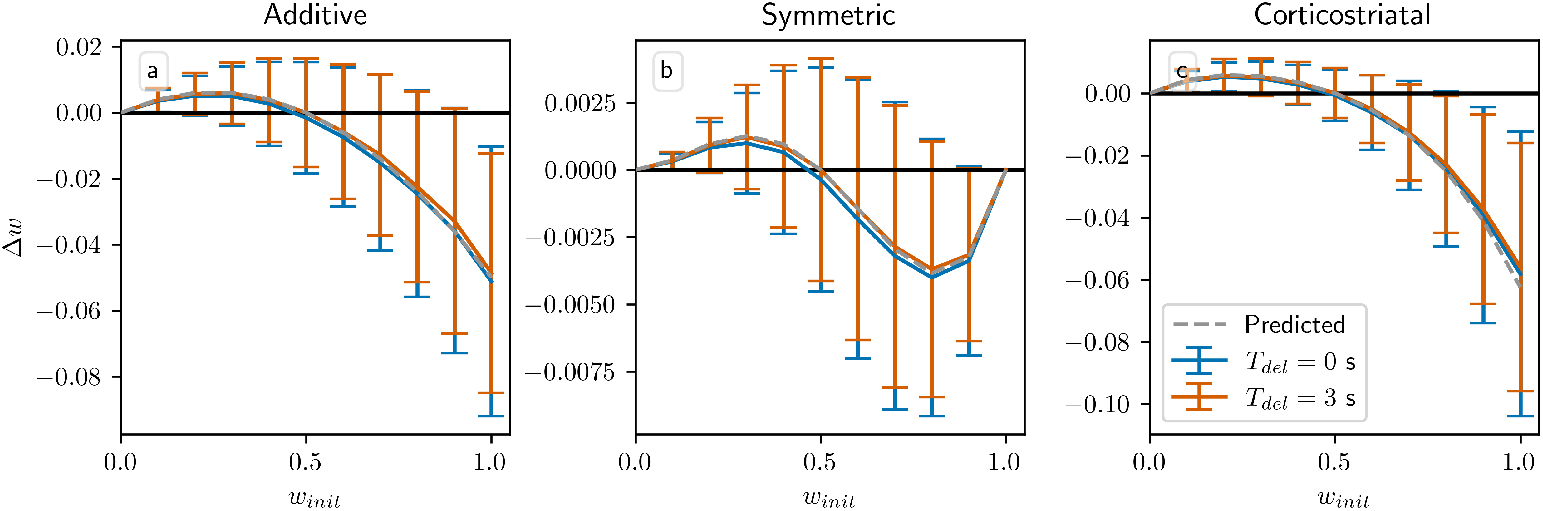
Weight drift after a single dopamine release in the value estimation setting with variable *T*_del_. Plots show results for the additive (a), symmetric (b), and corticostriatal (c) models, with *T*_del_ = 0 s and *T*_del_ = 3 s, as well as the predicted weight drift based on the averaged models, as *w*_init_ is varied for *N* = 1, with *p*_init_ = 0.5 fixed. Note that when *w*_init_ = 1 there are some deviations from predictions even for *T*_del_ = 3 s due to boundary effects not taken into account by the averaged models. Error bars show standard deviations over 10000 samples

Another assumption in our analysis is that *ϵ*, the time between presynaptic spikes and any postsynaptic spikes they cause, is small relative to *τ*, the time constant of synaptic plasticity. Specifically, we assume following [24] that *e*^−*ϵ/τ*^ ≈ 1; using our default values of *ϵ* = 0.001 s and *τ* = 0.02 s, this quantity is *e*^−*ϵ/τ*^ = 0.95. In Figure 11 we show the result of increasing *ϵ* to 0.005 s, in which case *e*^−*ϵ/τ*^ = 0.78. The main effect of increasing *ϵ* is to reduce the magnitude of the changes in weight. *τ* defines the duration of the window of synaptic plasticity; as *ϵ* increases, pre- and postsynaptic spikes grow farther apart relative to *τ*, and so weight changes due to presynaptic spikes directly causing postsynaptic spikes, corresponding to the *f*_+_(*w*_*i*_)*w*_*i*_*r*_*i*_ term in equation (16) for the additive and symmetric models (the other terms correspond to spike pairs that are close together only by chance), are reduced by a factor of *e*^−*ϵ/τ*^. Overall, though, for realistic values of *ϵ* the differences between the two curves are small, and the qualitative behavior is largely independent of *ϵ*.

**Fig. 11.**
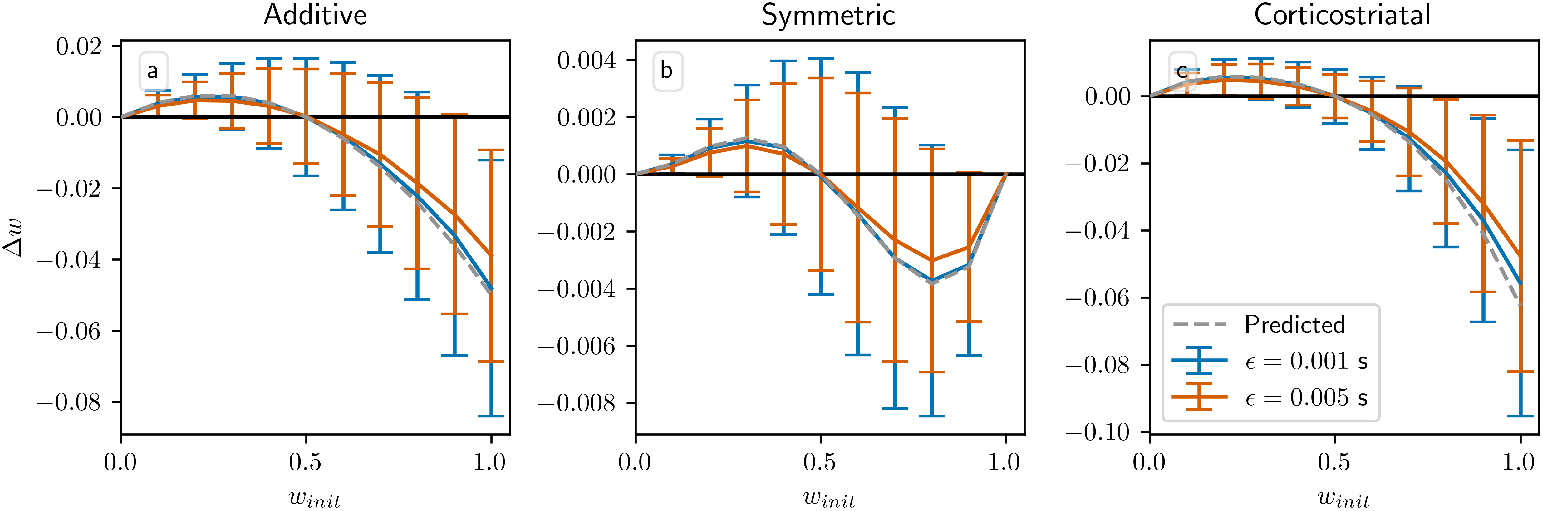
Weight drift after a single dopamine release in the value estimation setting across two choices of *ϵ*. Plots show results for the additive (a), symmetric (b), and corticostriatal (c) models, with *ϵ* = 0.001 s and *ϵ* = 0.005 s, as well as the predicted weight drift based on the averaged models, as *w*_init_ is varied for *N* = 1, with *p*_init_ = 0.5 fixed. Note that larger *ϵ* generally leads to less variability in Δ*w* and that when *w*_init_ = 1 there are some deviations from predictions even for *ϵ* = 0.001 s due to boundary effects not taken into account by the averaged models. Error bars show standard deviations over 10000 samples

## 4 Discussion

Accurately modeling learning in the cortico-basal ganglia-thalamic circuit requires the use of an appropriate synaptic weight update rule for the dopamine-dependent STDP in the corticostriatal connections. In this paper we examine three plasticity models that combine dopamine, eligibility, and spike timing signals in different ways – the additive, symmetric, and corticostriatal models – and evaluate their performance in various task settings. We find that the corticostriatal model is able to rapidly relearn swapped reward contingencies in the action selection setting. This rapid learning matches the results seen in experiments with animals [51] and humans [52]. However, the corticostriatal model performs poorly in the value estimation setting, as the synaptic weights do not converge to values that allow the agent to accurately estimate the rewards it will receive. In contrast, the additive and symmetric models are able to learn to accurately estimate reward values. But while they can accomplish a simple action selection task, they – especially the symmetric model – do not perform as well when reward contingencies occasionally switch, because their weights tend to become stuck at or near the boundaries of the domain. They are also more sensitive to *α* than the corticostriatal model in this setting, with larger values degrading their performance significantly. Overall, we find that the choice of which plasticity model to use can have a large impact on the dynamics of synaptic weights and hence on both the learning achieved by the circuit and the ability of the model to perform a given task. Which plasticity model is appropriate depends strongly on the tasks it will be asked to perform. Ultimately, these results suggest that different synaptic plasticity mechanisms may be at play at corticostriatal synapses involving different regions of the striatum with distinct functions, as well as at corticocortical synapses with dopamine-dependent plasticity [53], and that additional experimental and theoretical work is needed to pin down the precise forms of plasticity that occur at corticostriatal synapses and how they should be modeled.

Our mathematical analysis shows how the choice of parameter values impacts model performance on the tasks we considered. Specifically, we characterized how the stability of various equilibria (and therefore the ability of the models to solve each task) depends on the parameters *a*_sel_ (for the action selection setting), *α, τ, r*, 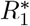, and 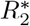. The parameters *α*, which represents the weighting of negative eligibility relative to positive in the additive and symmetric models and the weighting of weight decreases relative to increases in the corticostriatal model, and *τ*, the STDP time constant, have a strong impact on model performance In general, increasing *α* or *τ* will reduce the ranges of input firing rates *r* and rewards 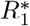, 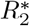 for which the system converges to the desired equilibria. Therefore, these parameters are particularly important for practitioners to understand and to select judiciously.

Why exactly do the three plasticity models run into difficulties in some settings? As discussed in Section 2.1.2, an important issue with the additive model is that it does not prevent weights from being driven to the boundaries (or past them, if the weights are not artificially cut off). The symmetric and corticostriatal models represent two alternative approaches to resolving this issue. (The multiplicative model used in prior work does not solve the issue when modified to incorporate dopamine.) The symmetric model keeps the weights bounded by multiplying the weight update equations by *w*(1 − *w*). However, in some cases this leads to the weights becoming stuck near the domain boundaries, resulting in poor performance in the action selection setting with contingency switching (Figure 5). The corticostriatal model, on the other hand, keeps the weights bounded by using scaling factors *w* or 1 − *w* depending on the sign of the product of the dopamine signal term with the eligibility trace term. In other words, it ensures that a scaling factor *w* is used when weights decrease and 1 − *w* when weights increase (see Table 2). The resulting model exhibits very good performance in the action selection setting with contingency switching. The cost of this plasticity rule, from an analytical perspective, is that the dopamine signal can no longer be factored out of the weight drift equation in the value estimation setting. Consequently the corticostriatal model features nonzero mean weight drift even when the mean dopamine signal is zero, leading to its failure to converge to the equilibria representing accurate value estimates in the value estimation setting.

Our modeling suggests a number of interesting hypotheses that could be tested experimentally. Most importantly, our findings suggest that different regions of the striatum may employ different plasticity mechanisms. For instance, the ventral and dorsal striatum may be specialized to their roles in value estimation and action selection, respectively [54]. We found that the additive and symmetric models are better suited to value estimation, while the corticostriatal model is better suited to action selection. This means that in regions of the brain involved in value estimation, we expect changes in synaptic strength to be roughly proportional to the average dopamine signal (difference from baseline), as seen in equation (16) describing the weight dynamics under the additive and symmetric models. Regions of the brain involved in action selection, meanwhile, will likely not display a clear proportional relationship between weight changes and dopamine signal, as the corticostriatal model lacks such a relationship (see equation (15)).

Another result that may be amenable to experimental validation is the finding that in many cases, stable learning of the correct behavior when using the additive or symmetric models requires that *τ* (*α* − 1) ∥*r*∥_1_ is small (see Theorem 1 and Table 4). This means that in regions where the additive or symmetric models accurately describe learning, we would expect the number of synaptic inputs or the input firing rates – and hence ∥*r*∥_1_ – to be anti-correlated with *τ*, the timescale of synaptic plasticity, and *α*, the scaling factor when eligibility is negative. Finally, our experiments with contingency switching suggest that in regions where flexibility is needed, we do not expect synaptic weights to be near their extremes of maximal or minimal strength.

One important issue that we have highlighted throughout this work is the impact that delays have on the weight dynamics. In the action selection setting, we saw that in order for effective learning to take place when there is a substantial delay between when we estimate the postsynaptic firing rate and when dopamine is actually delivered, we need to sustain activity in the selected channel. Without this sustained activity, performance quickly degrades as the delay increases (see Figure 6). With sustained activity, though, performance is maintained or actually improves as the delay increases. Likewise, in the value estimation setting, we require a long delay to ensure convergence to the equilibria representing accurate value estimation. Without this delay, we cannot guarantee convergence to the solution plane due to correlations between terms (although in practice this usually does not substantially affect the results). A complicating factor that we have not addressed is that experimental results consistently show that dopamine release immediately upon pre-post spike pairing does *not* lead to a change in weight; rather, the dopamine must come some time after the spiking activity to effect significant synaptic changes [34, 49]. Moreover, dopamine is not released instantly, but rather takes some time to ramp up to its peak value [48]. These findings raise important questions about how to best understand and model delays within a synaptic plasticity framework. Although we considered both the dopamine concentration and the eligibility trace as jumping up immediately and then decaying exponentially, for the sake of analytical tractability and for consistency with prior computational work, an important extension of these results would be to represent these signals as slowly ramping up and then ramping down over time and to study how these more realistic time-courses interact with delays and what computational outcomes they produce.

Our results build on past literature on spike-timing dependent and corticostriatal synaptic plasticity. Our additive model is based on the plasticity rules described in [24], but our plasticity rule differs from theirs in that we incorporate dopaminergic modulation of the synaptic plasticity. The settings in which we evaluate the model also differ significantly from theirs: while they study conditions under which symmetry breaking in the weight distributions occurs and when the models can learn to represent correlations in a set of inputs, we focus on measuring performance in more complex settings associated with the functional roles of corticostriatal synapses within corticobasal ganglia-thalamic circuits. The corticostriatal model in this paper is based on the plasticity model used in [21] but differs from their model in a number of important ways. We make several simplifications to the model, including setting the scaling factors and time constants for pre- and postsynaptic spike traces equal to each other (in their notation, *τ*_*PRE*_ = *τ*_*POST*_ and Δ_*PRE*_ = Δ_*POST*_ = 1), as well as considering a single class of striatal neurons rather than taking into account the existence of multiple striatal neuron subpopulations with different plasticity properties. They also employ their plasticity model in a more biologically realistic setting, incorporating many components of the basal ganglia circuitry that we leave out. A conceptually interesting difference between our models is that they use a single eligibility trace, summing up both positive (corresponding to pre-before-post spike pairs) and negative (post-before-pre) contributions, while we use two different traces for the positive and negative components. The use of two traces is justified by experimental evidence suggesting that the brain uses two distinct eligibility traces for LTP and LTD [37]. Note, however, that the computational model introduced in [37] differs considerably from the models used here, as it does not use *αw* or 1 − *w* factors to rescale the positive and negative traces, instead simply adding them together without modification. We find in Appendix D that altering our models to employ a single eligibility trace leads to qualitatively similar results in most cases, although the single-trace models are much more difficult to analyze mathematically. A number of other three-factor plasticity rules, which we did not consider, have also been explored in the literature. One important model can be found in [55]; while our learning rules are generally built from two-factor rules modified to incorporate dopaminergic feedback, they derive their learning rule directly from gradient ascent applied to a reward signal. Another work modeling dopamine-dependent STDP is [56]. The plasticity rule in that work closely resembles our additive model. However, while we focus on the corticostriatal synapses and employ a simple setting consisting of a population of cortical neurons connected to a single striatal neuron, they instead use a mixed population of excitatory and inhibitory neurons with random connectivity meant to model part of a cortical column. The scenarios that they use to test their model also differ from ours. For a more detailed review of other work on three-factor plasticity rules, see [3, 4].

In both the action selection and value estimation settings that we have considered, rewards are deterministic, with the agent always receiving reward 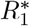 after taking action *A*_1_ and 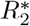 after taking *A*_2_. One could also consider versions of these settings with probabilistic rewards, where taking action *A*_1_ leads to reward 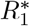 with some fixed probability *q* and to 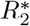 with probability 1 − *q*, and vice versa for action *A*_2_. In the value estimation setting, where we are able to calculate explicit formulas for the average dynamics (see Appendix A and the discussion in Section 3.2), it is easy to see that our results can accommodate probabilistic rewards by simply replacing 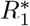 and 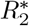 with 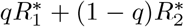 and 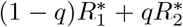, respectively, in the equations for the average dynamics (equations (16) and (17)). In the action selection setting the expressions for 𝔼[*R*\*] and 𝔼 [*p*] in equation (8) would need to be modified to properly compute the expected reward when rewards are probabilistic, but otherwise the qualitative behavior will be similar to the case of deterministic rewards. We also have not considered here how well synaptic weight values are preserved under dopamine release that is not associated with the modeled tasks but, as a diffuse signal, may reach the synapses of interest. A cost of the flexibility of the corticostriatal model, which works well in contingency switching, is a susceptibility to forgetting in such scenarios [57], which presumably might need to be countered by additional biological mechanisms.

Another direction for future work would be to extend our models to more complex settings in which both action probability learning and value learning are modeled in a biologically detailed way involving synaptic plasticity; possibly, thalamostriatal inputs, which we have ignored but have been considered elsewhere, could be important in this process as well [58, 59]. Notably, in our model and analysis, a hyperplane of equilibria provide accurate value estimation; this flexibility could allow for additional functionality, in that further constraints associated with more complicated task settings could be accommodated without compromising value estimation.

Future research could also examine whether our models can be adapted to more complex settings where exploration is necessary to achieve high reward. We generally focus on greedy reward maximization, in which we expect the probability of selecting the correct action to approach 1. Even in our contingency switching tasks there is no need to occasionally sample alternative actions, since we only swap rewards between the two actions: sampling one action gives complete information about the other as well. In other settings, for instance those where the rewards evolve independently, sampling alternative actions could lead to greater reward in the long term. The fact that the weights under the corticostriatal model do not converge to the extremal values of 0 or 1 in the action selection setting may be beneficial in such cases or others where exploration is useful. On the other hand, as described in Section 3.1.1, the limiting values of the weights under the corticostriatal model do not depend on the rewards or any other features of the environment, so more responsive exploration strategies would require additional circuitry to implement.

In addition to considering more complicated task settings, future work could include additional biological realism into our modeling framework. It is certainly possible that the plasticity mechanism used in the corticostriatal synapses incorporates features that are not well-captured by any of the models considered here. It is also possible that the simplified models that we consider omit aspects of the computational structure of the basal ganglia that are crucial for functional performance. For instance, we do not model the competition between direct and indirect pathways through the basal ganglia, nor the differing effects of dopamine on the two pathways [29, 32]. There may also be more complexity to dopaminergic feedback than the simple model we use; for example, recent work suggests that the dopamine signal may be better modeled as multidimensional rather than scalar-valued [60]. An exciting future direction would be to extend our analysis to take more of these subtleties into account. Nevertheless, we believe that the settings that we studied capture key computational components that will remain relevant in more detailed models and hence that our results will be informative in such settings.

Overall, what are the implications of our findings for models of the basal ganglia? We showed that each model does well in some settings yet fails to accomplish the given task in others. An interesting possible implication of our work is that different regions of the striatum may feature different plasticity mechanisms specialized to their particular roles. For instance, the ventral and dorsal striatum, which primarily contribute to value estimation and action selection, respectively [54], may use distinct plasticity rules tuned to the specific tasks that they perform, as suggested by experimental evidence [15, 17]. More generally, while we have focused in this paper on the corticostriatal connections, our framework is general enough that it may apply to any other region of the brain that receives dopaminergic signals, such as the prefrontal cortex, where dopamine-dependent plasticity also occurs [53]. The task settings we considered represent fairly general models of different aspects of the learning dynamics under dopaminergic feedback that do not depend heavily on the architecture of the basal ganglia. Thus, the fact that no plasticity rule performed well in every setting in our study supports the general principle that different brain regions should exhibit features – in this case, synaptic plasticity mechanisms – that are specialized for the specific computational functions that they perform.

## Acknowledgements

The authors acknowledge support from National Institutes of Health awards R01DA059993 and R01DA053014 and National Science Foundation award DMS-1951095. We thank Timothy Verstynen of Carnegie Mellon University for comments on an earlier draft of this manuscript and all members of the exploratory intelligence group for their feedback.

## Appendix A

**Averaged Model, Value Estimation Setting**

### A.1 Additive and Symmetric Models

Here we derive an averaged weight change model that adds up all contributions of pre-post and post-pre spike pairs and takes the average over realizations of the pre- and postsynaptic spike trains and over the possible timings of the dopamine signal, focusing on the additive and symmetric models in the value estimation setting. The presentation here largely follows that in [24], but with dopamine now included. (Much of the analysis also applies to the multiplicative model described in Section 2.1, but we do not focus on it here. For an analysis of the multiplicative model in a related setting, see [57].) We first give an expression for the total change in weight induced by a single triplet of a presynaptic spike at *t*_pre_, a postsynaptic spike at *t*_post_, and a dopamine signal *D* at *t*_dop_. This can be found by integrating over the time since the largest of *t*_pre_, *t*_post_, and *t*_dop_, because prior to *t*_pre_ or *t*_post_, the eligibility trace is zero, and prior to *t*_dop_, the dopamine trace is zero. The result for the additive and symmetric models takes the form

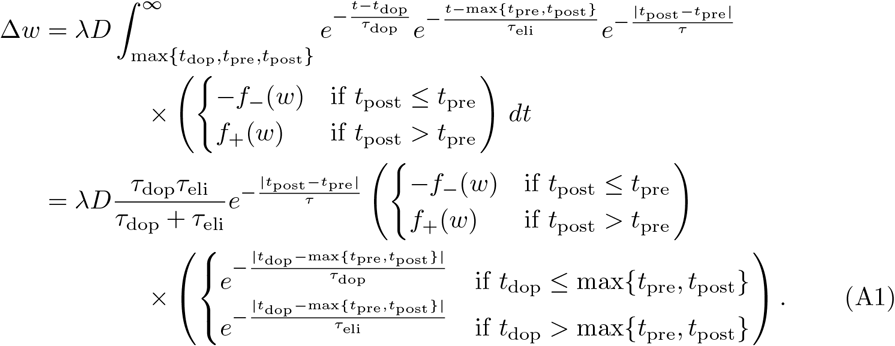

We restate here the definition of the dopamine signal for the value estimation setting:

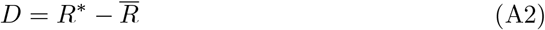

where

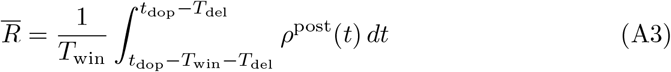

and *R*\* depends on the choice of action.

Following [24], we will define the cross-correlation functions 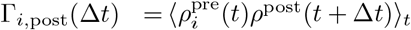, where ⟨·⟩_*t*_ denotes averaging over time. These will arise in the averaging process. We also define the point process *ρ*^dop^ indicating when a dopamine signal is delivered, with rate ⟨*ρ*^dop^(*t*) ⟩_*t*_ = *r*^dop^. (As noted previously, in simulations we assume for simplicity that dopamine is delivered periodically, but the precise form of the dopamine process does not matter as long as it has the given mean rate, it is independent of the spike trains, and dopamine signals are far enough apart that their interactions can be neglected.) Treating Δ*w* as a function of *t*_pre_, *t*_post_, and *t*_dop_, we can write the mean weight drift as follows:

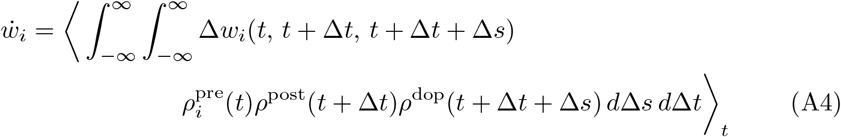

where *t* = *t*_pre_, Δ*t* = *t*_post_ − *t*_pre_, and Δ*s* = *t*_dop_ − *t*_post_.

Note that *ρ*^dop^ is independent of the other terms. Also, if *T*_del_ is large enough, we can assume that 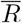 (and hence *D*) is independent of *ρ*^post^ (and hence also of 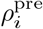, as 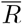 only depends on 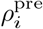 through *ρ*^post^), because any postsynaptic spikes counted by the integral in equation (A3) must occur at least *T*_del_ before the dopamine signal, and consequently, either the negative exponential in |*t*_dop_ − max{*t*_pre_, *t*_post_}| or the one in |*t*_post_ − *t*_pre_| in equation (A1) will be very small. Thus, assuming *D* is independent of the other terms provides a very good approximation if *T*_del_ is large enough. Another simplifying assumption we will make is that the weights change only a small amount on each dopamine release, so that we can treat *w*_*i*_ as constant in these expressions. Under these assumptions, we can substitute equations (A1) to (A3) into equation (A4) and split it into the *t*_post_ ≤ *t*_pre_ and *t*_post_ > *t*_pre_ cases as follows:

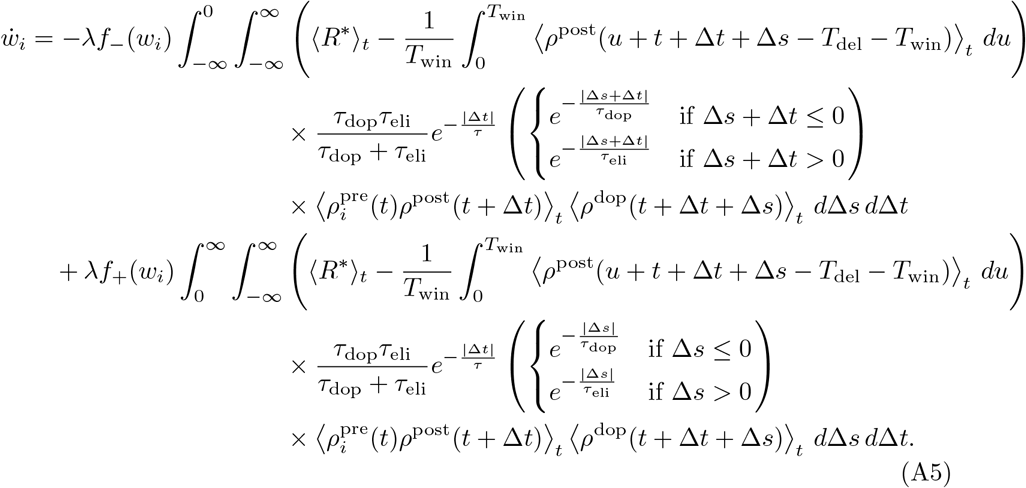

We can write ⟨*R*\*⟩_*t*_ as

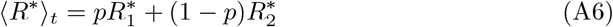

where *p* denotes the probability of selecting action *A*_1_. We will derive dynamics for *p* below; for now we simply assume that, like *w*, it changes only a small amount on each dopamine release and so can be treated as a constant here.

Recall that the postsynaptic firing rate is given by

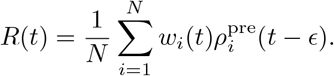

It follows that for any *x*,

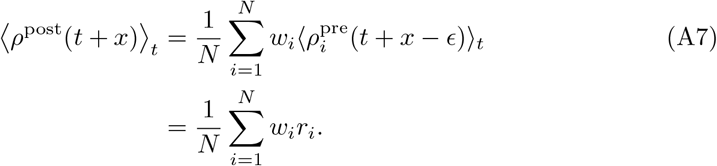

In particular equation (A7) applies to ⟨*ρ*^post^(*u* + *t* + Δ*t* + Δ*s* − *T*_del_ − *T*_win_)⟩_*t*_ in equation (A5). Additionally, ⟨*ρ*^dop^(*t* + Δ*t* + Δ*s*)⟩_*t*_ = *r*^dop^ is a constant. We can therefore make the change of variables Δ*s* ← Δ*s* +Δ*t* to combine the positive and negative integrals, arriving at the formula:

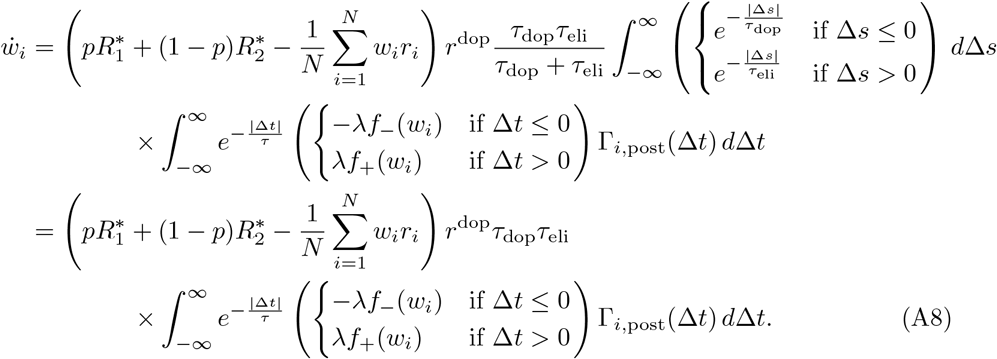

Note that the remaining integral is exactly the one found in [24]. Using equation (A7) and following [24], we decompose Γ_*i*,post_ as

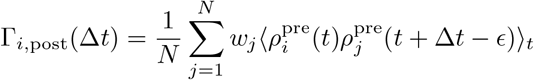

and define the normalized cross-correlation function

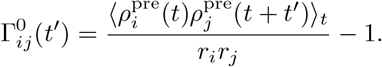

(Note that [24] assumes all presynaptic firing rates are identical, and so uses *r*^2^ in the denominator instead.) We also define the effective cross-correlation matrices *C*^±^ with elements

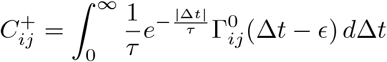

and similarly for 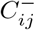 (which integrates from −∞ to 0). Then we can rewrite the integrals in terms of 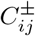 :

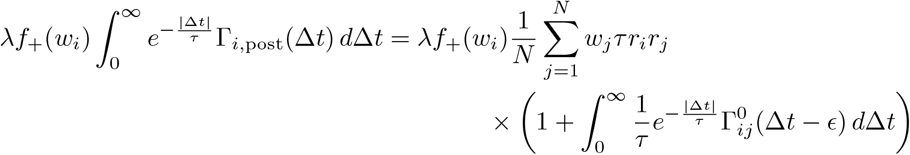

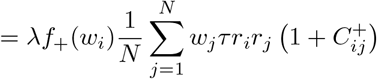

and similarly for the negative terms. Like in [24], we assume 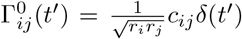 for some constants *c*_*ij*_ ≥ 0 (again extending their formula to non-identical presynaptic firing rates). Since the argument of 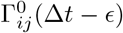 is never zero when Δ*t <* 0, it follows that 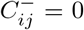 and 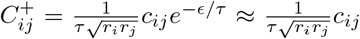. (We assume, as in [24], that *ϵ* is small enough that *e*^−*ϵ/τ*^ ≈ 1.) For Possion spike trains, the constants *c*_*ij*_ equal 1 if the spike trains are identical (because the autocorrelation is ⟨*ρ*(*t*)*ρ*(*t* + *t*^′^)⟩_*t*_ = *r*^2^ + *rδ*(*t*^′^) for a Poisson spike train *ρ* with rate *r*) and are otherwise less than 1. We will assume that the presynaptic spike trains are uncorrelated, so *c*_*ij*_ = 0 for *i* ≠ *j*. Therefore the formulas simplify as follows:

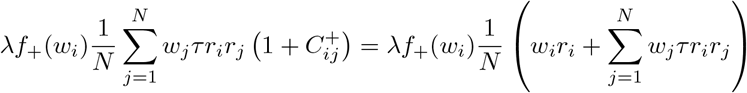

and

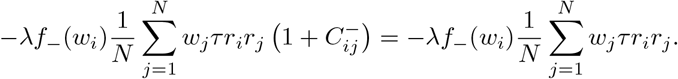

Substituting these results back into equation (A8), we obtain the formula for 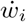:

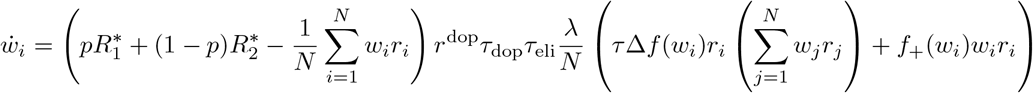

where Δ*f* = (*f*_+_ − *f*_−_). In vector notation, this formula can be written as:

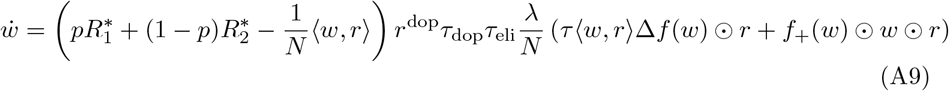

where ⊙ is the entrywise or Hadamard product and we treat *f*_±_(*w*) as applying entrywise.

We now derive dynamics for *p*, the probability of picking action *A*_1_. Recall that *π*_1_ and *π*_2_ obey

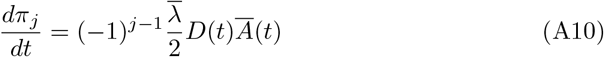

where 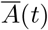 indicates the previous action selected, 1 for *A*_1_ and −1 for *A*_2_. The change in *π*_*j*_ for a single dopamine release is then given by:

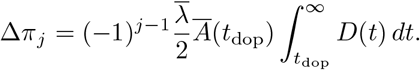

Taking an average only at the times dopamine is released, we see that

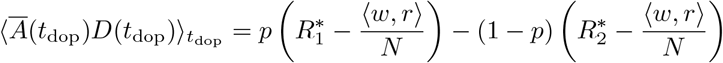

using equation (A6) and equation (A7) as above, assuming that *p* and *w* change only a small amount on each dopamine release. Incorporating the exponential decay of the dopamine signal over time, we get the average drift equation

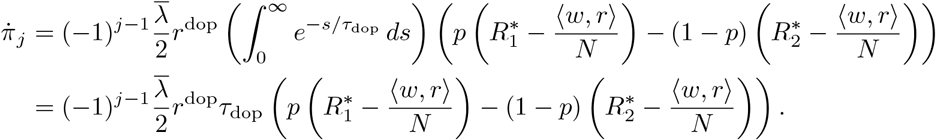

We can use this to derive an equation for *p*. From equation (10), we know that

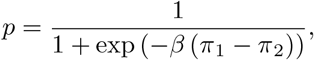

the standard logistic function applied to the difference *β* (*π*_1_ − *π*_2_). It follows that

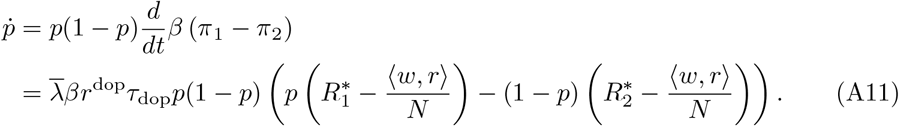

Thus we end up with an *N* + 1-dimensional system of differential equations for the *N* components of *w* along with *p*, given by equations (A9) and (17).

### A.2 Corticostriatal Model

The analogous expression to equation (A1) for the corticostriatal model is:

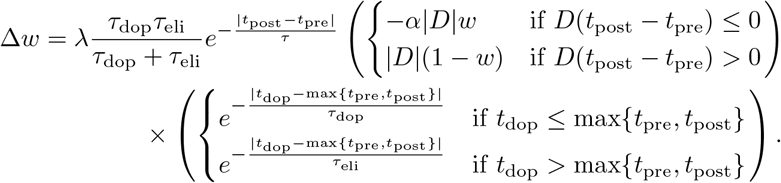

To derive an averaged form of the corticostriatal model, we need to decompose the expected dopamine signal into 𝔼 [*D*] = *D*_+_ + *D*_−_, where

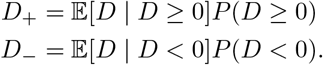

These can be computed by counting the number of postsynaptic spikes to fall inside the window in equation (A3), using the cumulative distribution function of the Poisson distribution and averaging over the two actions; on the *D* ≥ 0 side,

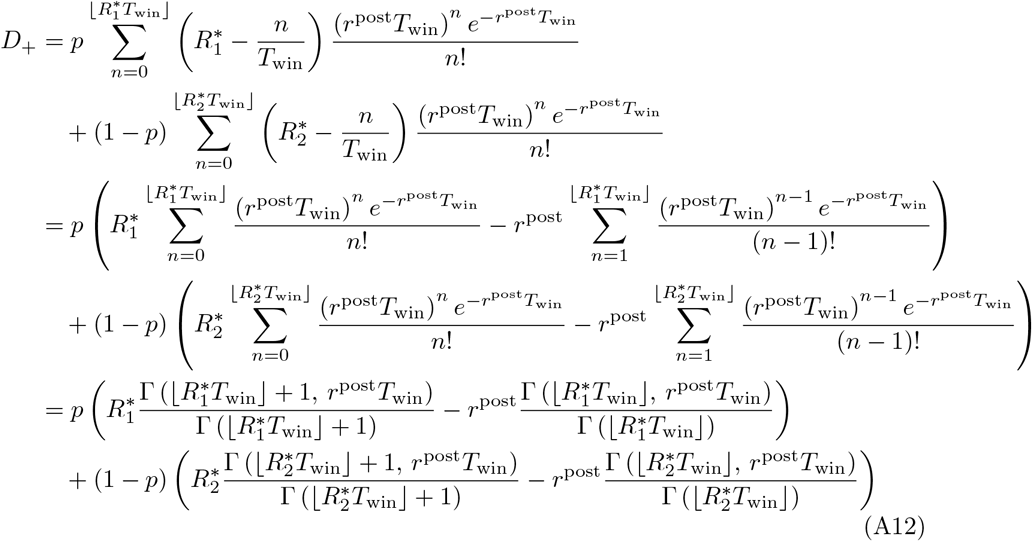

where *r*^post^ = ⟨*w, r*⟩*/N* is the postsynaptic firing rate. Since 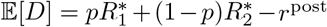, it follows that 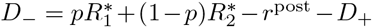. Then an analogous derivation to that in Appendix A.1, treating the *D* ≥ 0 and *D <* 0 cases separately, gives the following average weight drift formula

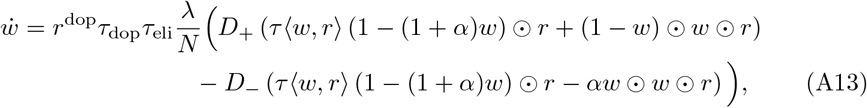

with 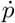 obeying equation (A11).

## Appendix B

**Analysis of Equilibria, Value Estimation Setting**

### B.1 Equilibria with *p* = 0 or *p* = 1

It is clear from equations (A9) and (17) that the following sets of points are equilibria for the additive and symmetric models:

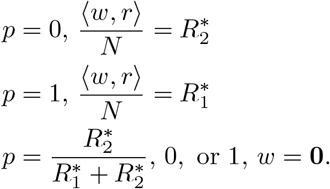

We now analyze the first two sets of points, which define *N* − 1-dimensional hyperplanes of weights. Assuming that 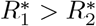, so that action *A*_1_ is more rewarding than *A*_2_, we would like to find conditions such that *p* converges to 1 and *w* converges to a point on the hyperplane 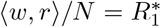, indicating that the system is correctly predicting the rewards it is receiving. (In the following analysis we generally assume for simplicity that each *r*_*i*_ is nonzero, since the *w*_*i*_ coordinate becomes trivial if *r*_*i*_ = 0.)

#### Theorem 2

*Pick w and p so that either p* = 1, 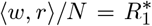 *or p* = 0, 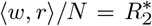. *For the additive and symmetric models given by equations* (A9) *and* (17), *the Jacobian at this equilibrium has one nonzero eigenvalue, which has an eigenvector that is zero in the p coordinate. This eigenvalue is negative if and only if*

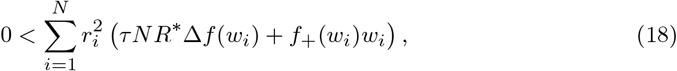

*with the choice of* 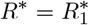 *or* 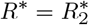 *corresponding to the equilibrium under examination*.

*Proof* The Jacobian at these points can be computed as follows:

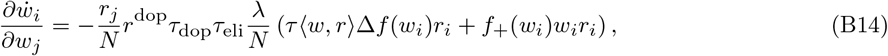

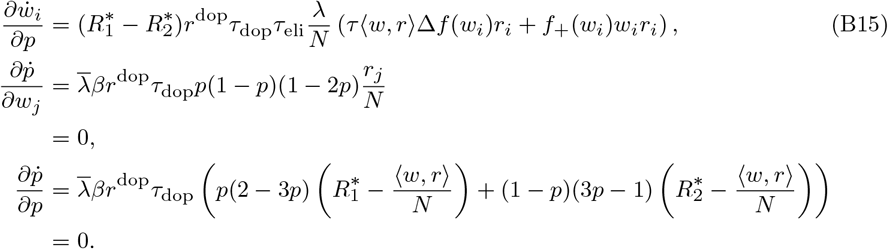

We will write the Jacobian matrix as

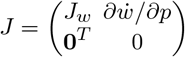

where

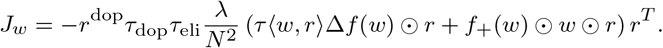

The submatrix *J*_*w*_ has eigenvalue 0 with multiplicity *N* −1 corresponding to the subspace orthogonal to *r*, that is, parallel to the planes 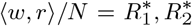. Its remaining eigenvalue is given by

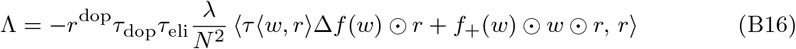

with associated eigenvector

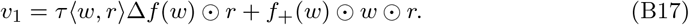

In general, the condition Λ < 0 (indicating stability) will define a subset of the planes, the shape of which depends on the precise form of *f*_+_ and *f*_−_:

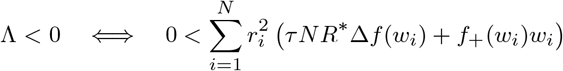

where we have used the fact that ⟨*w, r*⟩ */N* = *R*\* on the plane (with the choice of 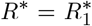 or 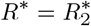 depending on which equilibrium we are examining).

The full matrix *J* retains this eigenvalue (the associated eigenvector simply has 0 in the *p* dimension). The eigenvalue 0 has multiplicity *N* for *J*. Like *J*_*w*_, it has *N* − 1 eigenvectors orthogonal to *r* (with 0 in the *p* dimension) and additionally has the eigenvector

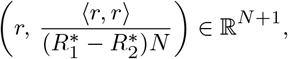

which is parallel to line segments stretching from the *p* = 0 hyperplane to the closest point on the *p* = 1 hyperplane.

To summarize, at *p* = 0 and *p* = 1 there exist hyperplanes of equilibria defined by 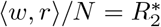 and 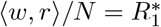, respectively. The Jacobian on these planes has one nonzero eigenvalue, Λ, which determines the direction of flow off the plane in the *w* direction (with zero component in the *p* direction). For the additive model, *f*_+_ = 1 and *f*_−_ = *α*, so the stability condition found above can be rewritten as follows:

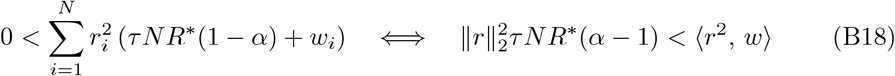

where *r*^2^ is the vector consisting of elements 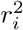. This condition holds on a half-space with a boundary perpendicular to *r*^2^ which will generally divide the hyperplanes of equilibria into two domains, one stable and one unstable. Note that if *α* = 1 this condition is always true for *w*_*i*_ ∈ (0, 1], so the entire plane is stable in this direction; however, in general we may have *α* > 1, in which case this global stability will not hold.

For the symmetric model, *f*_+_(*w*) = *w*(1 − *w*) and *f*_−_ = *αw*(1 − *w*). The stability condition is therefore a cubic polynomial in *w*:

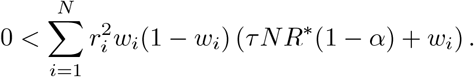

Like with the additive model, this condition is always satisfied when *α* = 1 (unless *w*_*i*_ = 0 or *w*_*i*_ = 1 for all *i*). In general, though, the region where this condition is satisfied will be more complex.

At each of these equilibria there also exists a center eigenvector with a nonzero component in the *p* coordinate, which we call the nontrivial center manifold. We now examine the behavior of the system along the nontrivial center manifolds:

#### Theorem 3

*Points of the form p* = 0, 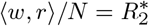 *or p* = 1, 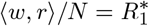 *are equilibria of equations* (A9) *and* (17) *with a nontrivial center manifold; the flow along this manifold is towards p* = 1 *and away from p* = 0 *if* 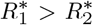, *or away from p* = 1 *and towards p* = 0 *if* 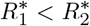.

*Proof* We focus for now on the *p* = 0 case. Let *w*_0_ be a point that satisfies 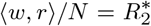, so that (*w*_0_, 0) is an equilibrium of equations (A9) and (17), and let 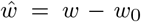. Let *v*_1_ ∈ ℝ^*N*^ be the eigenvector of *J*_*w*_ at the point (*w*_0_, 0) defined in equation (B17) (associated with the single nonzero eigenvalue), let *v*_2_, …, *v*_*N*_ be vectors orthogonal to *r* (so that they span the zero eigenspace of *J*_*w*_), and assume without loss of generality that ⟨*v*_*i*_, *v*_*j*_ ⟩ = *δ*_*ij*_ for *i, j* = 2, …, *N* so that the vectors are orthonormal. Let *V*_1_ ∈ ℝ^*N* ×*N*^ be the matrix with columns *v*_1_, *v*_2_, …, *v*_*N*_. We can then use the following coordinate transformation to align the axes of the system with the eigenvectors of *J* :

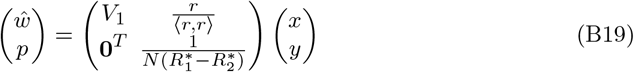

where *x* ∈ ℝ^*N*^ and *y* is a scalar multiple of *p*. Let *V* ∈ ℝ ^(*N* +1)×(*N* +1)^ be the entire coordinate transform matrix.

The only nonzero eigenvalue is associated with the coordinate *x*_1_, so the center manifold is of the form *x*_1_ = *h*(*x*_2…*N*_, *y*) (where *x*_2…*N*_ = (*x*_2_, *x*_3_, …, *x*_*N*_) ∈ ℝ^*N* −1^) such that *h*(**0**) = 0 and ∇*h*(**0**) = **0**. It follows that on the manifold,

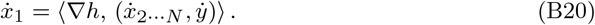

First note that using equation (B19),

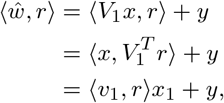

and by taking the time derivative we get

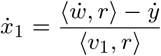

(since 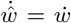). To address the right-hand-side of equation (B20), we introduce the *N* × *N* matrix

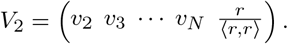

Since *V*_2_ is orthonormal, we have

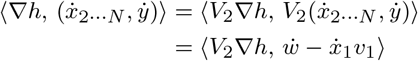

by taking the time derivative of the 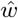 coordinates of equation (B19). Combining these results, we obtain

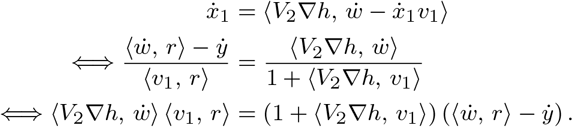

Note that ∇*h* is first order, so the terms involving 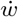 or 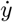 multiplied by ∇*h* are strictly higher order than the terms involving only 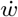 or 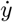 and can be dropped, leaving:

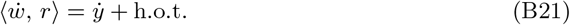

(Here “h.o.t.” stands for higher-order terms.)

We now collect the leading-order terms in this expression. To see the behavior of 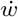 near the equilibrium (*w, p*) = (*w*_0_, 0), where we recall that 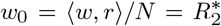, we substitute 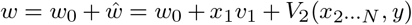 and 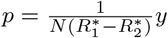 into equation (A9) to compute

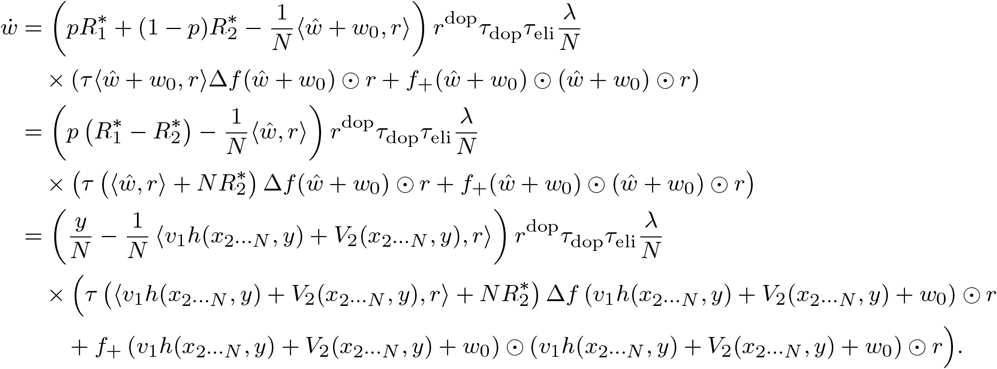

We know that ⟨*V*_2_(*x*_2…*N*_, *y*), *r*⟩ = *y*, so this expression can be written

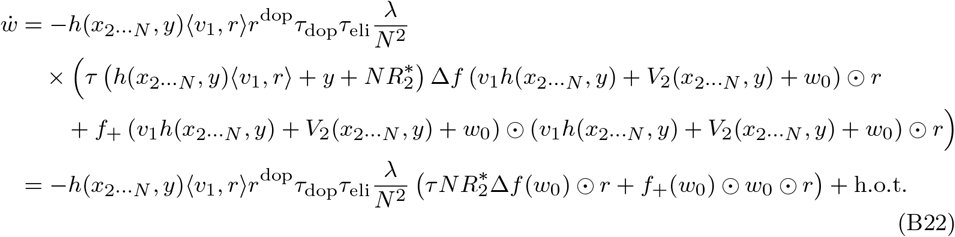

Similarly,

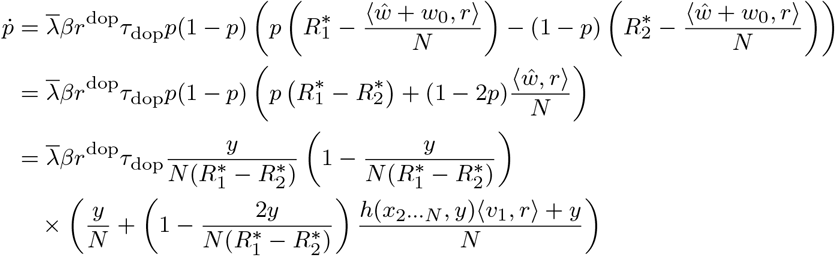

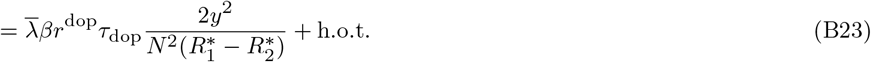

Moreover, equation (B23) implies that

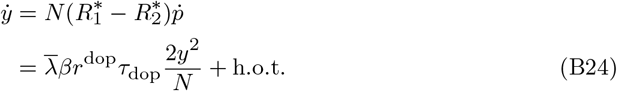

We can use equations (B22) and (B24) in conjunction with equation (B21) to derive an expression for *h*:

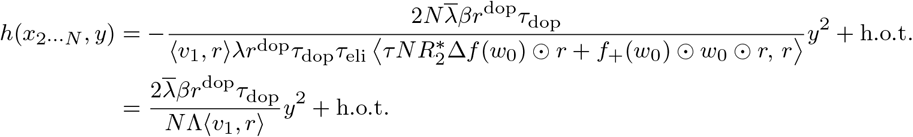

using the definitions of *v*_1_ and Λ (equations (B16) and (B17)); we will abbreviate this as *h*(*x*_2…*N*_, *y*) = *cy*^2^ + h.o.t. Note that *c <* 0 because Λ has the opposite sign as ⟨*v*_1_, *r*⟩.

Translating back into (*w, p*) coordinates, this means that we can parameterize the nontrivial center manifold near the equilibrium (*w*_0_, 0) as follows:

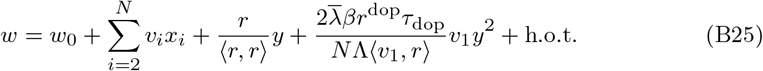

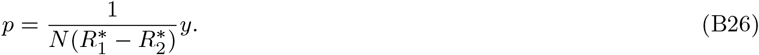

That is, the manifold consists of a linear term along the vector *r*, a quadratic term along the vector −*v*_1_ (negative since Λ has the opposite sign as ⟨*v*_1_, *r*⟩), and arbitrary translations orthogonal to *r* (from the *v*_2_, …, *v*_*N*_ terms).

We can now determine the dynamics on the manifold near the equilibrium. First, the flow in the *y* direction is always positive by equation (B24). Since 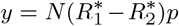 and *p* ≥ 0, the sign of 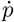 is determined by the sign of 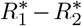 if 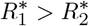, then 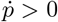, implying that the flow is away from *p* = 0; if 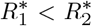, then 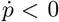, implying that the flow is towards *p* = 0. Thus the flow along this manifold promotes selection of the action that leads to the larger reward.

Meanwhile, the flow in *w* is described by equation (B22), which can be rewritten as follows using the definitions of Λ and *v*_1_:

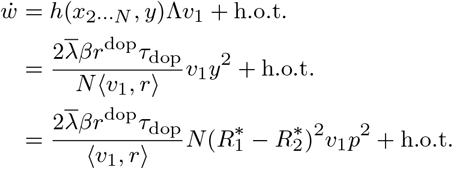

If ⟨*v*_1_, *r*⟩ > 0, then Λ < 0, so that trajectories near the equilibrium converge to it exponentially in the *w* axes, while on the manifold, there is a slow flow in the direction of *v*_1_. If ⟨*v*_1_, *r*⟩ < 0, then trajectories diverge exponentially from the equilibrium, while on the manifold, the slow flow is in the direction of −*v*_1_. In the *N* = 1 case shown in Figure 7, the sign of ⟨*v*_1_, *r*⟩ simply equals the sign of the scalar *v*_1_, so trajectories along the center manifold always increase in *w* when sufficiently close to the equilibrium.

The symmetry of the system makes analysis of the center manifold at the *p* = 1 equilibrium very similar to that of the *p* = 0 case. In fact, replacing *p* with 1 −*p* only has the effect of swapping the roles of 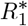 and 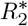. Concretely, we select *w*_0_ such that 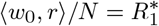 (so that the point *w* = *w*_0_ lies on the hyperplane of equilibria associated with *p* = 1), let 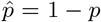, and make the change of coordinates 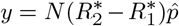, with the order of 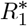 and 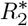 reversed. It can then be seen that the formula to parameterize the manifold near (*w, p*) = (*w*_0_, 1) is identical to equation (B25) for *w* (although *w*_0_ is different, and Λ and *v*_1_ also depend on the choice of *w*_0_), while the expression for *p* is

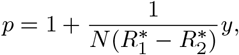

where in this case we will have *y <* 0 to parameterize values of *p* in [0, 1]. Similarly, the sign of 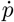 will be positive if 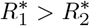, but in this case that implies the flow will be *towards p* = 1, while 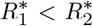 implies the flow will be away from *p* = 1: in both cases the flow is towards picking the action that leads to the larger reward.

### B.2 Equilibria with *w* = 0

As mentioned previously, in addition to the equilibria at *p* = 0 and *p* = 1 there are also equilibria at *w* = **0** and 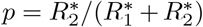, 0, or 1. The Jacobian at each of these points depends on the choice of *f*_+_ and *f*_−_. We first analyze the additive model, where *f*_+_ = 1 and *f*_−_ = *α*.

#### Proposition 4

*For the additive model, if the condition*

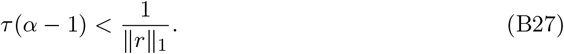

*holds, then the w* = **0**, *p* = 0 *and w* = **0**, *p* = 1 *equilibria are saddles where the Jacobian has exactly one negative eigenvalue (corresponding to the eigenvector* (0, …, 0, 1)*), with all other eigenvalues real and positive. They gain at least one zero or negative eigenvalue if the condition does not hold. The w* = **0**, 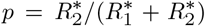 *equilibrium has a Jacobian with only positive real eigenvalues if equation* (B27) *holds, but gains at least one zero or negative eigenvalue if the condition fails*.

*Proof* The Jacobian can be computed as follows:

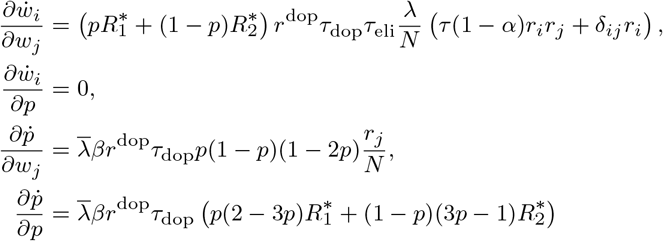

for *p* = 0, 1, 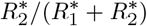. We will write the Jacobian matrix as

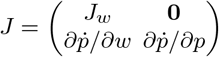

where

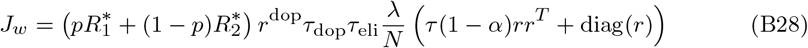

is symmetric.

As *J* is block-triangular, one eigenvalue is given by 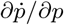, which varies depending on which equilibrium we are considering. The corresponding eigenvector is (0, …, 0, 1) ∈ ℝ^*N* +1^, pointing entirely along the *p* axis. The remainder of the eigenvalues are eigenvalues of *J*_*w*_. We now show that *J*_*w*_ is positive definite if and only if the condition given in equation (B27) holds.

To see that this condition determines the definiteness of *J*_*w*_, we examine ⟨*x, J*_*w*_*x*⟩ for *x* ∈ ℝ^*N*^ :

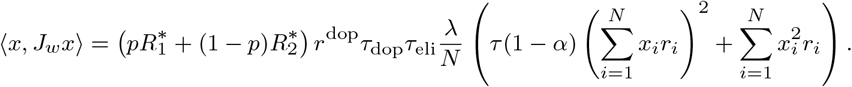

We assume *r*_*i*_ > 0 for each *i*, which allows us to apply Sedrakyan’s inequality^2^ [61]:

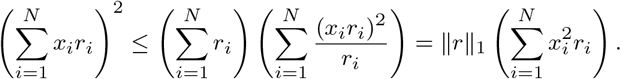

Suppose that 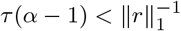. Then we have the bound:

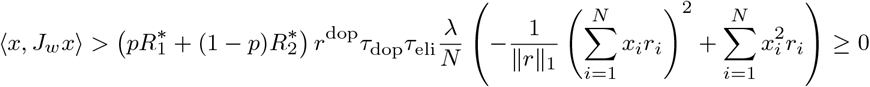

by Sedrakyan’s inequality, so *J*_*w*_ is positive definite. Conversely, if ⟨*x, J*_*w*_*x*⟩ > 0 for all *x*, then in particular it must hold for *x* = **1**, so we get:

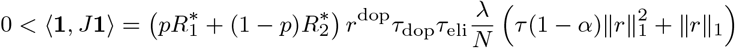

which can be rearranged to give 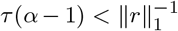. Thus we see that if equation (B27) holds, *J*_*w*_ only has positive eigenvalues, whereas if it does not hold, it must have at least one zero or negative eigenvalue.

The overall stability of the equilibria also depend on 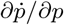 as follows:

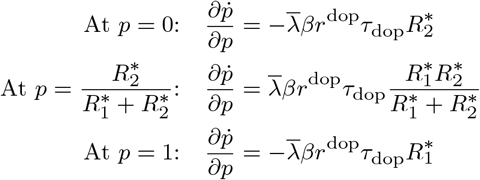

So the *w* = **0**, *p* = 0 and *w* = **0**, *p* = 1 equilibria are saddles with exactly one negative eigenvalue (corresponding to the eigenvector (0, …, 0, 1)) if equation (B27) holds, and they gain at least one zero or negative eigenvalue if the condition does not hold. The *w* = **0**, 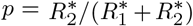 equilibrium is unstable (i.e. only positive eigenvalues) if equation (B27) holds, but it gains some zero or negative eigenvalues if the condition fails.

As a useful special case, when *N* = 1, *J* has only two eigenvalues, so one eigenvalue is determined by 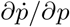 while the sign of the second eigenvalue is determined by equation (B27): it is positive when the condition holds and zero or negative when it fails. When *N* > 1, we cannot determine exactly how many eigenvalues change sign when the condition fails, only that at least one must change sign. We do not analyze the eigenspaces at these equilibria further because we were unable to find closed-form solutions for the eigenvalues and eigenvectors of *J*_*w*_ for arbitrary *r*.

Note that the condition given in equation (B27) is equivalent to checking whether Λ > 0 using equation (B18) at the point *w* = **1***NR*\**/* ∥*r*∥_1_ (for 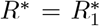 or 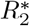). In other words, if this condition holds, then the point at which the vector **1** intersects the hyperplanes of equilibria analyzed in Appendix B.1 is unstable, Λ > 0. Conversely, if this condition fails to hold, then the point at which **1** intersects the hyperplanes has Λ ≤ 0.

For the symmetric model, *f*_+_(*w*) = *w*(1−*w*) and *f*_−_(*w*) = *αw*(1−*w*). Consequently, at *w* = **0**, *J*_*w*_ is the zero matrix, so the zero eigenvalue of *J* has multiplicity *N*. The final eigenvalue, with eigenvector (0, …, 0, 1), is equal to 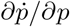, which is identical to the expressions computed above for the additive model: at *p* = 0 and *p* = 1 it is negative, while at 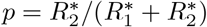 it is positive.

We do not fully analyze the zero eigenspaces at these critical points. However, note that the equation for 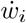 for the symmetric model is identical to the corresponding equation for the additive model multiplied by *w*_*i*_(1 − *w*_*i*_), which does not change the sign. This means that the qualitative behavior of the system – towards or away from these equilibria – will be the same as in the case of the additive model, although the dynamics here are higher-order in *w* due to the *w*_*i*_(1 − *w*_*i*_) term.

### B.3 Extra Equilibria in the Symmetric Model

The symmetric model has a number of additional equilibria not present in the additive model. There are equilibria at *w* = **1** and either *p* = 0, 1, or 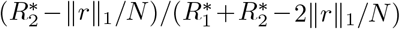. More generally, for any partition of the index set *I* ∪ *J* ∪ *K* = {1, 2, …, *N*}, there are equilibria at points *w* of the form

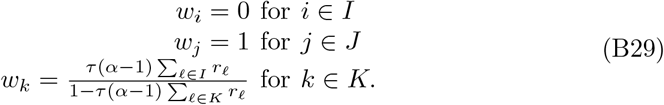

This requires that 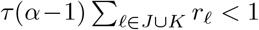 to ensure that 0 *< w*_*k*_ < 1 if *K* is nontrivial. These points are equilibria when either *p* = 0, *p* = 1, or

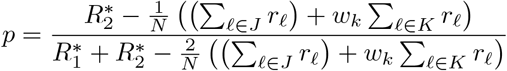

where *w*_*k*_ is as above. We will focus our analysis only on the *w* = **1** equilibria, as most of our experiments employ *N* = 1 and the other equilibria require *N* > 1.

#### Proposition 5

*Suppose* 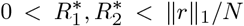. *If equation* (B27) *holds, then under the symmetric model the w* = **1**, *p* = 0 *and w* = **1**, *p* = 1 *equilibria have all positive eigenvalues, whereas if* 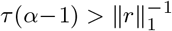, *then they are saddles with one positive eigenvalue and N negative eigenvalues. Meanwhile, the w* = **1**, 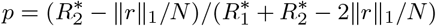 *equilibrium is a saddle with one negative eigenvalue and N positive eigenvalues if equation* (B27) *holds and if* 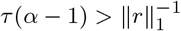, *then it is stable*.

*Proof* The Jacobian at these equilibria can be computed as follows:

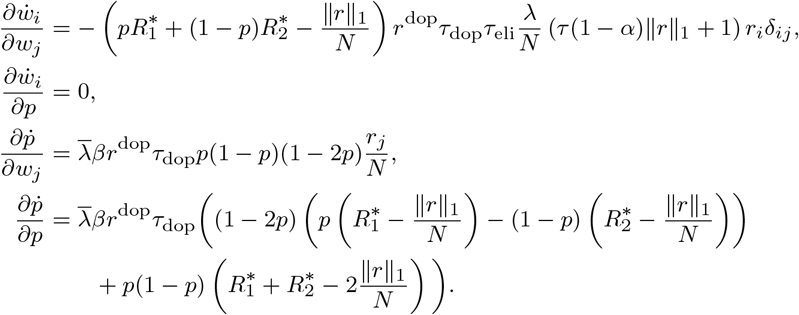

Write the Jacobian matrix as

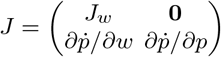

where

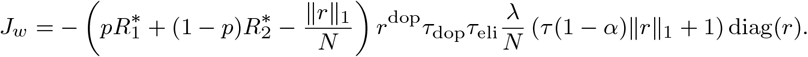

The eigenvalues of *J*_*w*_ are multiples of the entries of *r*. We typically assume *r*_*i*_ > 0 for all *i*, and for these fixed points we assume 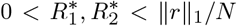. (If 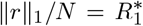 or 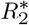 is allowed, *J*_*w*_ will be zero at either *p* = 0 or *p* = 1; for simplicity, we do not examine this case.) Recall that *α* ≥ 1. The signs of the eigenvalues are therefore determined by the same condition as that we found for the additive model, equation (B27): if 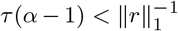, then all eigenvalues of *J*_*w*_ are positive, while if 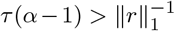, then all eigenvalues are negative.

The last eigenvalue (with eigenvector (0, …, 0, 1)) equals 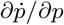:

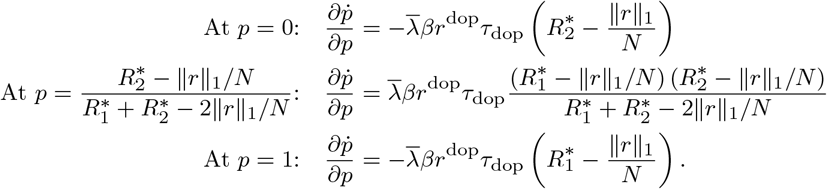

Since 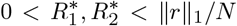, the first and third of these quantities are positive, while the second is negative. This, combined with our analysis of *J*_*w*_, gives the stability conditions described above.

### B.4 Corticostriatal Model

Since *D*_+_ and *D*_−_ depend on an incomplete gamma function applied to *w*, it is infeasible to analytically solve for most of the equilibria of the system when using the corticostriatal model. We can, however, identify the equilibria at *w* = **0**, because at this point 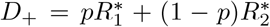 (see equation (A12)) and so *D*_−_ = 0. It follows that, as in the additive and symmetric models, *w* = **0**, *p* = 0, 1, and 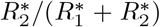 are equilibria. The Jacobian at these points can be computed as follows:

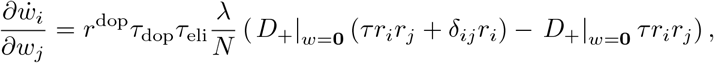

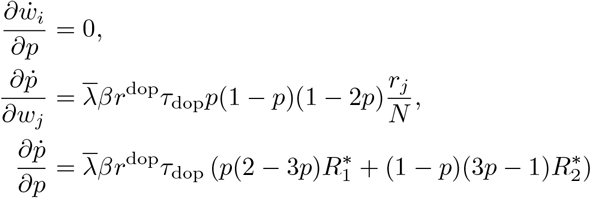

for *p* = 0, 1, 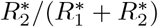. At *w* = **0**, 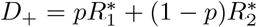 and *D*_−_ = 0, so *J*_*w*_ is given by:

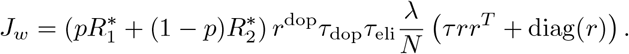

All other terms are the same as in Appendix B.2. It is easy to see that *J*_*w*_ is positive definite, as diag(*r*) is positive definite (since *r*_*i*_ > 0 for all *i*) and *rr*^*T*^ is positive semidefinite, so their sum must be positive definite. We can calculate 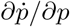 as in Appendix B.2 to get that the *w* = **0**, *p* = 0 and *w* = **0**, *p* = 1 equilibria have Jacobians with one only one negative eigenvalue and the rest positive, while for the *w* = **0**, 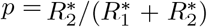 equilibrium, the Jacobian has only positive eigenvalues, so only the unstable eigenspace is nontrivial.

As can be seen from Figure 7, however, the corticostriatal model also has additional equilibria at nonzero values of *w*, for which we do not have closed-form expressions.

## Appendix C

**Action Selection Setting**

### C.1 Averaged Model

Our derivation of the averaged model for the action selection setting resembles the derivation in Appendix A, with some important modifications. Recall that in this setting, an action is selected in the window [*t*_dop_ − *T*_del_ − *T*_win_, *t*_dop_ − *T*_del_], by measuring the relative activity in each channel as defined in equation (A3), and then spiking activity in the non-selected channel is completely suppressed until the next action selection window, while activity in the selected channel is reduced by a factor of *a*_sel_. To model this scenario, we need to separately consider the cases where action *A*_1_ or *A*_2_ is selected. Using equation (8), we have:

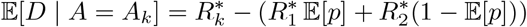

and the probability of picking action *A*_*k*_, when averaged over instantiations of the spike trains, is given by 𝔼 [*p*] or 1 − 𝔼 [*p*] for *k* = 1 or 2, respectively, as defined in equation (9). When computing 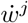, we only include the 𝔼 [*D* | *A* = *A*_*j*_] term, since when *k* ≠ *j* all activity is suppressed, and we replace *r* with the rescaled firing rate vector *a*_sel_*r*, since activity in the selected channel is reduced by this factor. (Note that we use the original *r* in the computation of 𝔼 [*p*], since activity is not suppressed during the action selection window.) Finally, we assume, as we did in the value estimation setting, that *T*_del_ is large, so that activity in the action selection window will be approximately independent of the eligibility values of the weights at the later time when dopamine is released.

With the modifications described above, an analogous calculation to that made in the value estimation setting gives the following equation for the average weight drift for the additive and symmetric models:

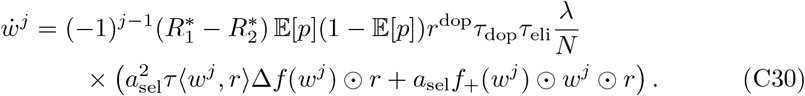

For the corticostriatal model, we must take into account that the form of the weight update equation depends on the sign of the dopamine signal as described in equation (5). If 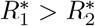, as we typically assume, then the dopamine signal will always be greater than or equal to zero when action *A*_1_ is selected, and less than or equal to zero when *A*_2_ is selected. Since channel 1 is only active when *A*_1_ is selected and is suppressed when *A*_2_ is selected, the equation for 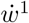 only includes the terms for positive dopamine, and likewise the equation for 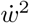 only includes the terms for negative dopamine. We therefore obtain the following system of equations:

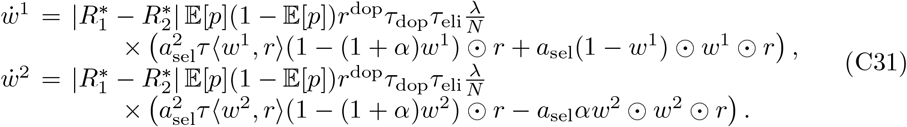

Note that each term in these equations has the same sign as the corresponding term in equation (A13) because *D*_+_ ≥ 0 and *D*_−_ ≤ 0. If 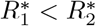, then we simply swap the equations.

### C.2 Dynamics

For both the additive and symmetric models in the action selection setting, each 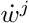 has a zero at *w*^*j*^ = **0**. The symmetric model has zeros at *w*^*j*^ = **1**, due to the 1 − *w* terms in *f*_+_, *f*_−_, as well as the combinations of these points: *w*^1^ = **0**, *w*^2^ = **1** and *w*^1^ = **1**, *w*^2^ = **0**. Additionally, when *N* > 1 each 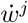 for the symmetric model has zeros at the same points found in the value estimation case, specified in equation (B29); we do not analyze these points further.

In general, we cannot directly compute the Jacobians at these equilibria because of the 𝔼 [*p*] terms, which depend on *w* via equation (7). However, it is still possible to describe the dynamics in many cases. We now prove the theorem stated in the main text:

#### Theorem 1

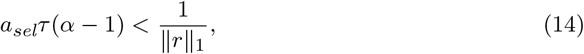

*and suppose* 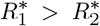. *Then the w*^1^ = *w*^2^ = **0** *equilibrium is a saddle that is repelling along the w*^1^ *coordinates and attracting along the w*^2^ *coordinates. For the symmetric model, w*^1^ = **1**, *w*^2^ = **0** *is an attractor, w*^1^ = **0**, *w*^2^ = **1** *is a repelling equilibrium, and w*^1^ = *w*^2^ = **1** *is a saddle that is attracting along the w*^1^ *coordinates and repelling along the w*^2^ *coordinates*.

*Proof* We will determine the dynamics by studying the simplified system,

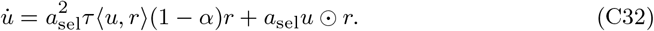

Since 𝔼 [*p*] > 0, the dynamical equations for 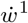 and 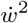 for both additive and symmetric models (equation (C30)) are equal to substituting *w* for *u* in equation (C32) and multiplying by a function that is either always greater than or equal to zero or always less than or equal to zero, depending on the sign of 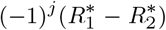. It follows that any zero of 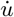 is also an equilibrium of the original system, and that the direction of flow near an equilibrium is either identical or reversed, again depending on the sign of 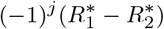.

The Jacobian of this system at *u* = **0** is given by:

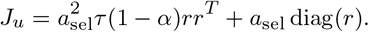

This formula closely resembles equation (B28), the formula for *J*_*w*_ in Appendix B.2. Calculations analogous to those performed there show that *J*_*u*_ is positive definite if and only if equation (14) holds.

It follows that for both the additive and symmetric models, if equation (14) holds and 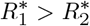, then near the equilibrium *w*^1^ = *w*^2^ = **0** we will have 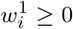 and 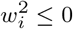 for all *i*.

To determine the stability of the symmetric model at *w*^*j*^ = **1**, we use the system

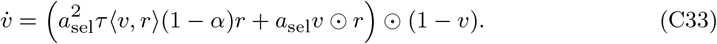

Its Jacobian at *v* = **1** is:

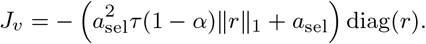

The eigenvalues of *J*_*v*_ are multiples of the entries of *r*, and it can readily be seen that if equation (14) holds (and *r*_*i*_ > 0 for all *i*, as we generally assume), then *J*_*v*_ is negative definite.

It follows that if equation (14) holds and 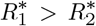, then the point **1** is attracting for *w*^1^ and repelling for *w*^2^. Combining these results, we see that under the symmetric model, *w*^1^ = **1**, *w*^2^ = **0** is an attractor, *w*^1^ = **0**, *w*^2^ = **1** is a repelling equilibrium, and *w*^1^ = *w*^2^ = **1** is a saddle which is attracting along the *w*^1^ coordinates and repelling along the *w*^2^ coordinates.

For the additive model, this leads to relatively simple dynamics. If 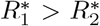, *w*^1^ will generally grow until it hits the boundary at **1**, while *w*^2^ will approach the origin. For the symmetric model, the dynamics are similar when *N* = 1, except that **1** is a genuine zero of 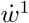, rather than just an accumulation point caused by clipping the weights to [0, 1]. When 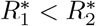 all these relations are reversed; thus in both settings the dynamics promote selecting the more rewarding action. When *N* > 1, though, the dynamics under the symmetric model may be more complex due to the presence of other equilibria.

For the corticostriatal model, in addition to the zeros at *w*^*j*^ = **0**, there is also a set of zeros that differ for the *w*^1^ and *w*^2^ equations. Assuming 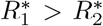, so that equation (C31) is used, the points *w*^1*^ and *w*^2*^ defined below are zeros of 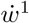 and 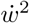, respectively:

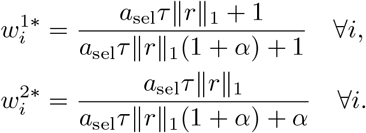

We cannot directly compute the Jacobians at these points because of the 𝔼 [*p*] terms. However, we can determine their stability by considering the reduced system

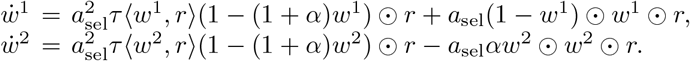

This system is produced by dividing the terms in equation (C31) by strictly positive functions, so it has the same equilibria with identical stability properties. The Jacobians at 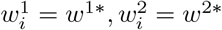 are:

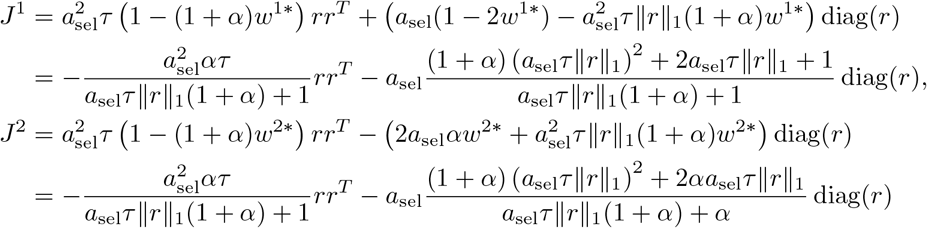

which are both negative definite matrices. It follows that both *w*^1*^ and *w*^2*^ are attractors for their respective equations, so that *w*^1^ converges to *w*^1*^ and *w*^2^ converges to *w*^2*^. Since *w*^1*^ > *w*^2*^, the more rewarding action, *A*_1_, will be favored over *A*_2_. The 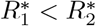 case can be treated similarly.

## Appendix D

**Single Eligibility Trace**

We now revisit the question of whether to use a single eligibility trace summing up both positive (corresponding to pre-before-post spike pairs) and negative (post-before-pre) contributions, as is done in [21], or to use two different traces for the positive and negative components, as we do elsewhere in the paper. We had several reasons for focusing on models with two different eligibility traces. One was analytical convenience: the use of two traces is necessitated by the assumption, made by [24, 26] as well as in this paper, that the contributions to the weight changes made by individual spike pairs sum independently, a natural assumption that greatly simplifies analysis. With only one eligibility trace, different spike pairs may cancel each other out, rendering this independence assumption invalid. We therefore cannot derive averaged forms of the single-trace models like we did for the two-trace models. A second justification for the focus on two-trace models is that there is experimental evidence suggesting that the brain in fact uses two different traces, one for LTP and one for LTD [37], at least in cortical pyramidal neurons.

Nonetheless, for modeling purposes, it is important to compare the performance of the single-trace model to the model two-trace model that we have considered. We therefore test a single-trace version of our model that replaces equation (3) with

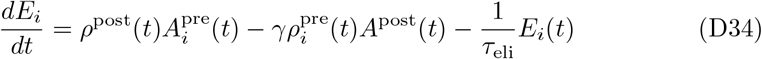

where *γ* ≥ 1 is a scaling parameter controlling the strength of negative eligibility terms relative to positive terms. The single-trace differential equation for the weights in the additive and symmetric cases is given by

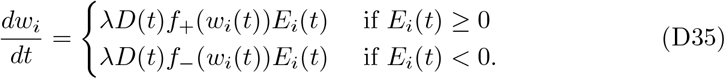

and for the corticostriatal model is given by

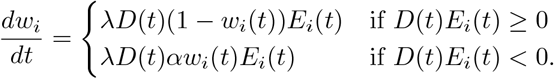

The single-trace version of the corticostriatal model is largely equivalent to the model described in [21], although they use different scaling factors and time constants for pre- and postsynaptic activity.

One important characteristic of the single-trace versions of the additive and symmetric models (equation (D35)) is that they are largely insensitive to variations in *α*. This insensitivity arises because *E*_*i*_(*t*) is usually positive, since presynaptic spikes directly cause postsynaptic spikes after a delay of *ϵ* and not vice versa, which tilts the balance to favor positive eligibility. Hence, the *α*-dependent *f*_−_ term is only rarely used. The *α* parameter is therefore not an effective way of adjusting the relative strengths of the positive and negative components of the eligibility trace. This observation motivates the introduction of the parameter *γ* in equation (D34) to provide a better means of controlling the relative strengths of the two components in the single-trace models. The single-trace corticostriatal model is still sensitive to *α* because it depends on the sign of the product *D*(*t*)*E*_*i*_(*t*), rather than just *E*_*i*_(*t*), so the term with *α* will have an impact when *E*_*i*_(*t*) > 0 and *D*(*t*) < 0.

We show simulations of the single-trace models in the action selection setting in Figures 12 and 13 (without and with contingency switching), and in the value estimation setting in Figures 14 and 15. In the action selection setting, we see that the behavior of the additive and symmetric models is qualitatively very similar to their behavior in the two-trace settings (compare Figure 12 to Figure 3 and Figure 13 to Figure 5), with varying *γ* having a very similar effect to varying *α* in the two-trace versions. The corticostriatal model also behaves similarly when *γ* is small, but as *γ* increases its behavior deviates significantly from that in the two-trace setting, which can lead to worse performance in some cases: the flow in Figure 12i is away from the optimal weight values *w*^1^ = 1, *w*^2^ = 0, while in Figure 3i it converges to (*w*^1*^, *w*^2*^) which, while not optimal, still leads to the correct action being selected.

**Fig. 12.**
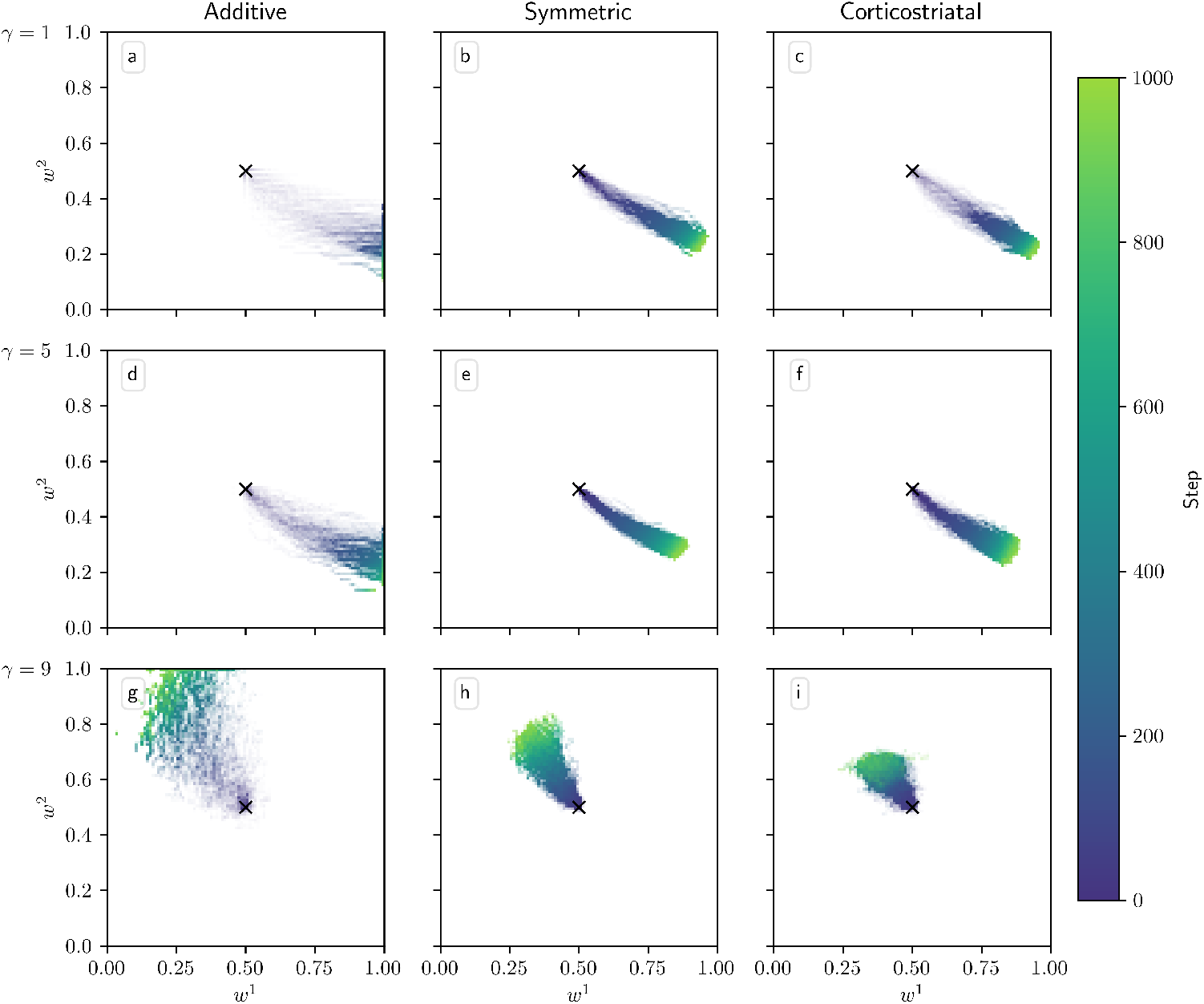
Distribution of *w*^1^ and *w*^2^ over time in the action selection setting for single-trace models as *γ* is varied. Columns show the additive (a, d, g), symmetric (b, e, h), and corticostriatal (c, f, i) models. *γ* is varied across rows: (a-c) *γ* = 1; (d-f) *γ* = 5; (g-i) *γ* = 9. We use *α* = 1 and *w*_init_ = 0.5 here, marked by the “×” in each plot. As we do not have averaged forms of the single-trace dynamics we do not include vector fields or fixed points here

**Fig. 13.**
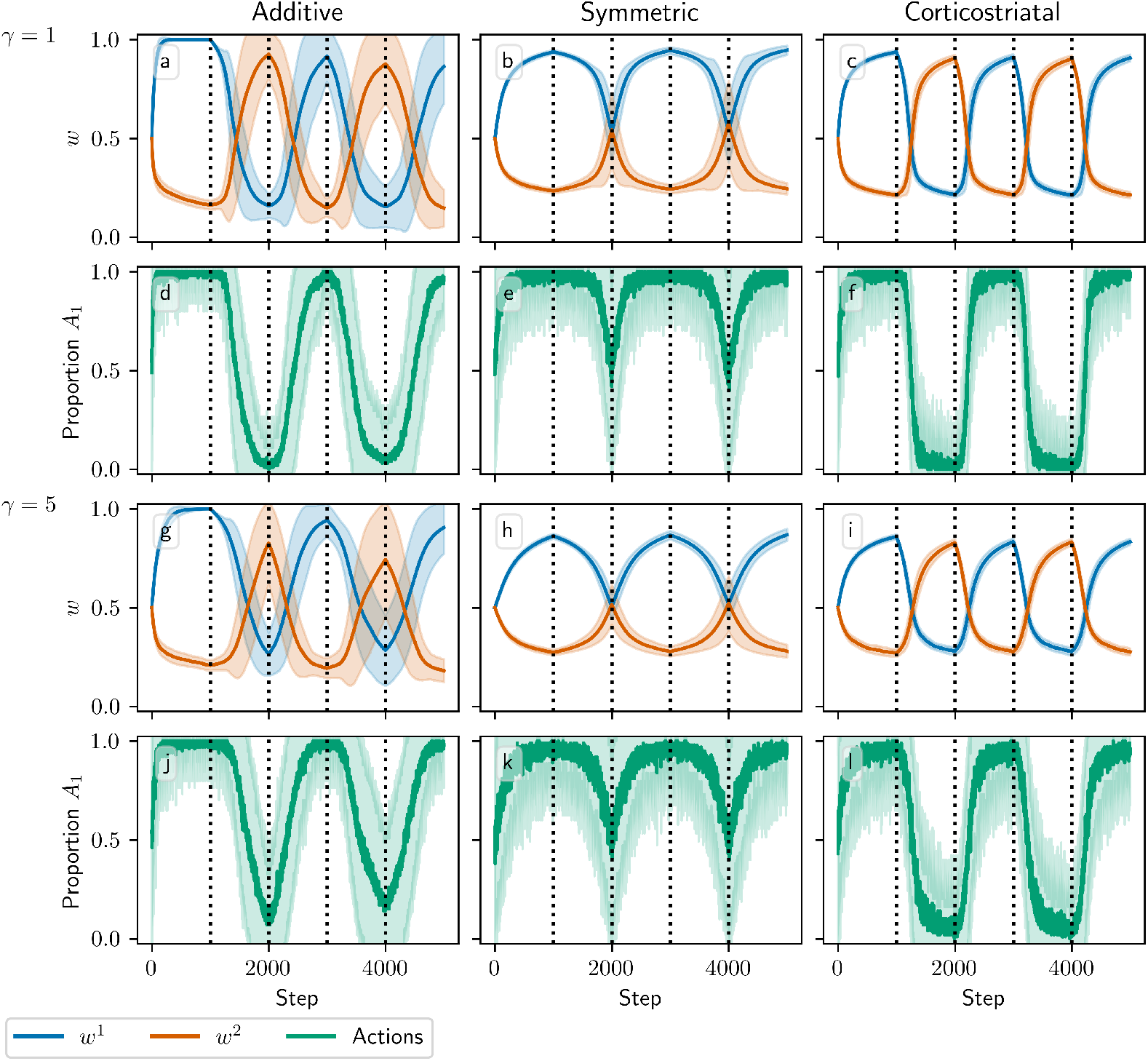
Model performance in the action selection setting with contingency switching for single-trace models as *γ* is varied. Plots show weights (a-c, g-i) and probability of taking the correct action (d-f, j-l) versus time for the additive (a, d, g, j), symmetric (b, e, h, k), and corticostriatal (c, f, i, l) models. *γ* is varied across rows: (a-f) *γ* = 1; (g-l) *γ* = 5. We use *α* = 1 and the other parameters are the same as those used in Figure 5

**Fig. 14.**
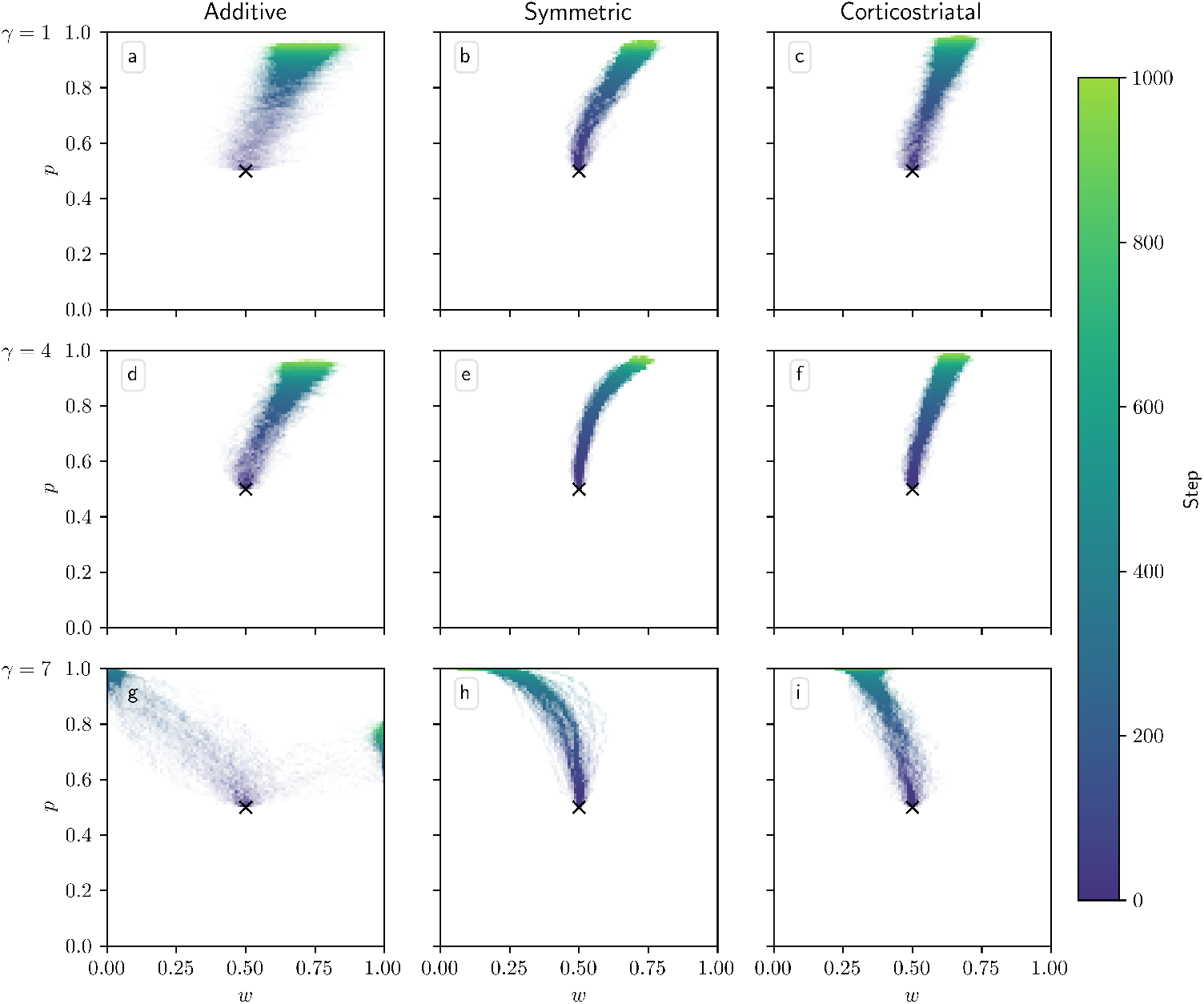
Distribution of weights over time in the value estimation setting for single-trace models as *γ* is varied. Columns show the additive (a, d, g), symmetric (b, e, h), and corticostriatal (c, f, i) models. *γ* is varied across rows: (a-c) *γ* = 1; (d-f) *γ* = 4; (g-i) *γ* = 7. We use *α* = 1 and *w*_init_ = 0.5 here, marked by the “×” in each plot. As we do not have averaged forms of the single-trace dynamics we do not include vector fields or fixed points here

**Fig. 15.**
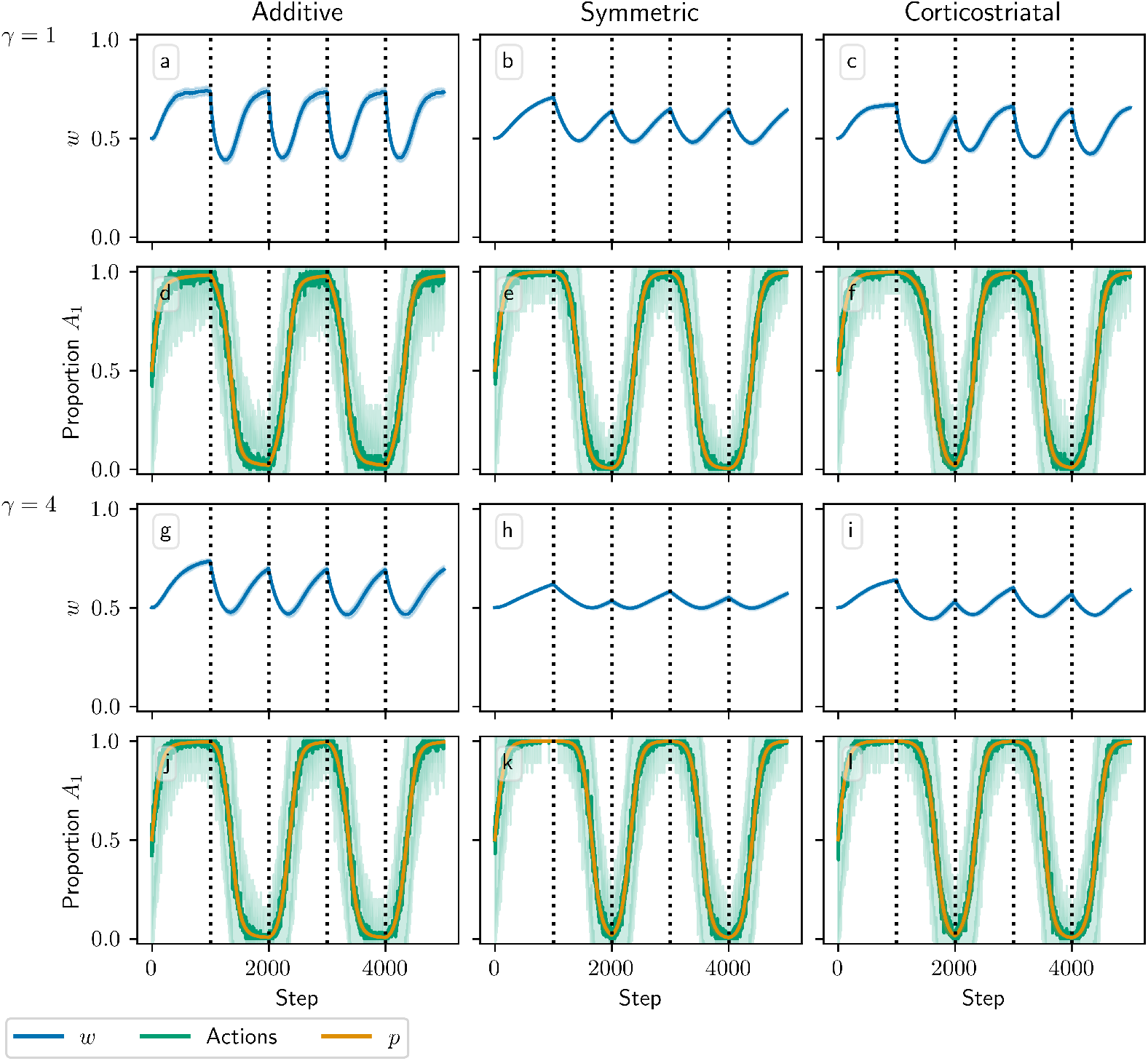
Model performance in the value estimation setting with contingency switching for single-trace models as *γ* is varied. Plots show weights (a-c, g-i) and the proportion of trials on which action *A*_1_ is picked along with the values of *p* (d-f, j-l) versus time for the additive (a, d, g, j), symmetric (b, e, h, k), and corticostriatal (c, f, i, l) models. *γ* is varied across rows: (a-f) *γ* = 1; (g-l) *γ* = 4. We use *α* = 1 and the other parameters are the same as those used in Figure 9

In the value estimation setting, we again see that the additive and symmetric models behave qualitatively similarly across the single-trace and two-trace versions of the dynamics (compare Figure 14 to Figure 7 and Figure 15 to Figure 9). Like in the action selection setting, as *γ* increases the corticostriatal model’s behavior deviates from that in the two-trace setting. In Figure 14f and i, this leads to convergence to somewhat different equilibria; in Figure 15i and l, this actually leads to improved performance over the two-trace version seen in Figure 9i and l (although as we noted previously, performance at this task is quite sensitive to the learning rates *λ* and 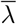, and better choices for their values may lead to better performance).

Overall, using a single eligibility trace does not seem to significantly improve performance in most cases, it makes the dynamics much more difficult to analyze, it complicates the weight equations for the additive and symmetric cases, and it requires tuning the artificial parameter *γ* to capture effects of *α* > 1. On the other hand, it does halve the number of eligibility trace equations that need to be simulated.

## Footnotes

1 We keep the additive model despite its problems because it provides a simple point of comparison; when we implement the model, we clip the weights to keep them between 0 and 1, which in theory could occur through some sort of biological mechanism that we do not consider here. With the weights thus constrained, the model in fact performs reasonably well in many cases.

2 Also known as Engel’s form or Titu’s lemma, it is a consequence of the Cauchy-Schwarz inequality.

